# Genome survival and descent in population pedigrees

**DOI:** 10.64898/2026.11.14.766719

**Authors:** Elizabeth Thompson

## Abstract

Motivated by issues of the maintenance of genetic variation in a highly endangered species, we continue earlier studies into the population genetics ancestry and descent of genome segments. In this paper, we focus on the survival and descent of genome in a population pedigree of a diploid dioecious species. We consider a population potentially being maintained from a small founder base, taking the canonical example a random-mating population of 100 males and 100 females. We show how previous methods to examine the ancestry of a single genome in such a population can also be used to account the survival genetic lengths and number of descendant copies of founder genome segments, and explore the variance of these measures. We show the variance of the population distribution over realizations within a pedigree is scarcely smaller then the variance over pedigrees: variation is in meiosis rather than in pedigree. While all founders continue to lose genome over time, the very first generations are critical to determining the amount of surviving genome. Dependence among founders is slight, except for between the two genomes of a founder: however when monogamy is enforced this high dependence extends to founder mates. We briefly consider extensions of population size (both smaller and larger), of mating structure, and of variance in offspring number, including variance in male reproductive success.

## 1 Introduction

In considering the maintenance of genetic diversity in a highly endangered diploid dioecious species, both ancestry and descent are relevant. The limited species genome ancestral to a small extant population defines what has survived, while descent from that population defines what may be maintained in future re-establishment of the species. We have previously focused on ancestry (Thompson, 2024), but we now continue the analysis of that paper, focusing on the survival and descent of genomes in small populations. The canonical example used will be a random-mating population of 100 males and 100 females each generation, although other population sizes and mating processes will also be considered.

For later convenience we introduce some notation. We consider a diploid dioecious population. At each generation there are *N* males and *N* females. There are thus 2*N* individuals, and 4*N* genomes at each generation. Simulations will be performed by genome (separating the maternal and paternal genome of each individual), but of course in scoring output the two genomes of any individual may be combined or compared. We consider a genome of genetic length *L* Morgans. This may be a full genome (e.g. *L* = 30 Morgans) and may or may not be divided into chromosomes (e.g. a genome of 30 autosomes each of length 1 Morgan (Thompson, 2023)). In other cases we consider only a part of an organism’s genome, for example *L* = 1 Morgan, but will still refer to this portion as a genome, to avoid confusion of multiple terms.

In studying the history of a small population, there will be a single known or unknown pedigree. In considering the potential future of the population, variation over realized pedigrees becomes the focus. We consider both variation due to realizations of meiosis in a single pedigree, and also variation among pedigrees resulting from the pedigree generating process. Most of the results presented are from simulations. While enough realizations are performed to be confident of comparisons, these are illustrative rather than definitively extensive.

In Section 2 we set the models and notation, and provide some historical background. We discuss how simulations of genome in a pedigree may be performed either forwards or backwards over the generations, We discuss simulation of ancestry of segments of a single haploid genome in a diploid dioecious population (Thompson, 2024), and then show how this same approach gives results for survival and descendant counts for segments of ancestral genomes to the current population. We conclude Section 2 with some basic results for unbounded populations that provide a basis for comparison with the small finite population of interest. Section 3 is the main results section. Throughout this section results are presented for a random-mating population with *N* = 100 males and *N* = 100 females in each generation. Population pedigrees are created by the random assignment of each of 100 sons and 100 daughters to their father and mother (the males and females of the preceding generation). In Section 3.1 we first consider the basic processes of extinction both of an individual’s. diploid descendants, and of their pair of haploid genomes, and then contrast patterns of ancestry and descent of diploid individuals. Section 3.2 considers descent of founder genome in more detail. For a single realized population over 100 generations and a 30 Morgan genome, we consider of the counts, lengths and distribution of segments of founder genome. We consider the relationship between descendants and genetic descendants, and between surviving genome and total descendant genome. In Section 3.3 we consider much more fully the variance of descent statistics, and the relationship of within-pedigree vs between-pedigree variation using the methods of Section 2.4. For this section only we focus on a 1 Morgan length of genome, such as a single chromosome, and on two rather different statistics: the proportions of surviving genome and the maximal number of descendant copies across the 1-Morgan genome. Within a single population pedigree chromosomes segregate independently, so the distribution of within-pedigree statistics can easily be extended to a broader genome. In Section 3.4 we consider dependence in the survival of founder genomes, showing how in a random mating population high dependence is only found between the two genomes of each diploid founder.

In Section 4 we briefly consider the effects of varying population size, mating structure and of variance in reproductive success.. Generally population size has little effect on extinction except in the smallest populations (Section 4.1.1), while a large population simply scales up counts of descendant genome segments (comparing Sections 3.2.1 and 4.1.2). Generally population size affects the time-scale of statistics such as rates of change of frequencies of specific founder genomes, but has little effect on events determined within a few generations, such as extinction and the dependence in descent of founder genomes. Mating structure can have a more dramatic effect on dependence among founder genomes (Section 4.2): in particular enforcing monogamy so all offspring fall into full sibships dramatically increases the dependence in descent from founder mates. Variance in reproductive success (Section 4.3) has a major effect on extinction, but apparently little effect on time-scale or segment descent conditional on survival. We conclude the paper with a brief discussion.

## 2 Descent and ancestry: models and methods

### 2.1 Pedigrees and meioses

In this subsection we establish some basic notation and terminology:

1. **The pedigree** First, there is the pedigree of the individuals, whether known from records or simulated under a random mating model or other population process. The pedigree defines the existence maternal and paternal links between parent and offspring individuals. A pedigree may be very compactly stored by specifying the mother and father of each non-founder individual., In simulating a random-mating population *N* males and *N* females over *k* generation, one simply assigns each of the 2*N* individuals at a given generation a father and a mother at the preceding one. Fathers (mothers) are a sample of size *N* sampled randomly with replacement from the males (females) of the parental generation;
2. **The genomes** Each individual has a maternal genome and a paternal genome, each of genetic length *L* Morgans, which may be divided into *C* chromosomes. (We consider only the autosomal genome in this paper.) The maternal (paternal) genome is determined by the individual’s maternal (paternal) meiosis, specifying the inheritance of DNA from his/her mother (father).
3. **The meioses:** The stochastic processes governing meiosis occur independently in every meiosis *j* of the population pedigree. The outcomes are most easily specified by meiosis indicators Thompson (1994), which, considering the genome as a continuum, in any meiosis *j* and at genome location *y* may be written as

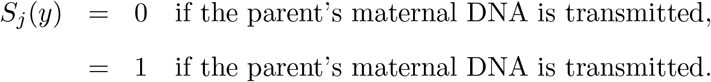

Under Mendel’s first law, the marginal distribution of *S_j_*(*y*) is given by

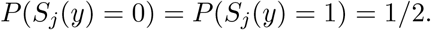

By the definition of genetic distance, the expected number of changes in *S_j_*(*y*) in any meiosis *j*, over a genetic length *y ∈* [0*, ℓ*] Morgans, is equal to *ℓ*.

A meiosis pattern on any chromosome may be very compactly stored by specifying *S_j_*(0), the value at the start of a chromosome, and the (normally few) locations *y* of changes in *S_j_*across the chromosome. We find that the division of genome *L* into *C* chromosomes has little effect on most results Thompson (2023). A chromosome break is simply a restart from the marginal 50/50 distribution of *S_j_*, and hence a 50/50 chance of a change. Not only are meioses *S_j_*(*y*) are independent over *j*, but they are Markov over *y* in the absence of genetic interference Haldane (1919). This is convenient for simulation and computation, but the Haldane model is not essential; any model may be used for the switch-points across a chromosome. For example, as noted by Thompson (2024), very small chromosomes may be badly modeled by a Haldane map function. For normal meiosis, at least one chiasma on the tetrad is required, leading to a minimum genetic length of 50 centiMorgans (cM), but in very small chromosomes a meiosis is very unlikely to have more than a single switch-point. Such chromosomes could be modeled with a probability 0.5 of no switches, and probability 0.5 of a single one at a random point (in terms of genetic distance).

Given pedigree and meiosis specifications, ancestry and descent is then a labelling process. Contiguous segments of DNA may be labelled with the haploid founder genome implied by descent from the ancestors. Following earlier papers both on pedigrees (Thompson, 2011) and in populations (Brown et al., 2012), we will denote these labeled haploid founder genomes as FGL. A genome segment may also be labeled by the set of current descendants to which it is ancestral. Labeling downwards by founder origin may seem simpler, since each point in every genome descends from a unique FGL, but may be ancestral to many current genomes. However, each point in every current genome traces to a unique genome at each preceding generation; tracing ancestry upwards is often more efficient even when downwards measures of genome survival and descent are the focus.

### 2.2 Simulation and computation on defined pedigrees

At a single locus, downward simulations have a long history. Edwards (1967) simulated single-locus descent in the pedigree of an entire small population to estimate the inbreeding coefficients of all extant individuals. Similar simulations were carried out in the pedigrees populations of severely endangered species (MacCluer et al., 1986) to estimate kinships, inbreeding, and founder allele survival. Interestingly, the first simulation for the estimating of inbreeding coefficients (Wright and McPhee, 1925) proceeded up rather than down the pedigree. The 50-50 meioses were realized up through the ancestors of an individual of interest, with careful specification of the order of realization and the checking of coancestry. The example was of a cattle pedigree in which there may be close inbreeding, and extensive cross-generational mating – for example a grandfather-granddaughter mating. The method does not seem to have been pursued, perhaps due to the practical difficulties of any simulation in 1925, with path-counting computations already established (Wright, 1922). However, upwards simulation of ancestry is quite practical in many contexts where phenotypic data do not complicate the process.

Observable data, allelic types, genotypes, or phenotypes have probabilities that are dependent on the identity by descent (IBD) patterns that are implied by the pedigree (1) and meioses (3), and the consequent labelling of Section 2.1. Data creates very complex constraints on the latent meioses, and with data typically available for the bottom generation(s) of the pedigree, it was natural that the first single-locus likelihood computation algorithms proceeded upwards and considered cases where information from the descendants of any individual could be independently accumulated to the individual’s parents (Elston and Stewart, 1971). However, computations can proceed in any direction through a large and complex pedigree, provided only that the requisite joint information accumulated on individuals is retained until it is no longer required (Cannings et al., 1978). Upwards simulation is usually infeasible in the presence of data. For example, in the case of ancestry of a rare recessive allele on a large complex pedigree (Geyer and Thompson, 1995), the rarity of the allele forces a likelihood that there are few founder copies. For a recessive lethal trait, an added constraint is that no ancestor can be homozygous for the allele (Thompson and Morgan, 1989) .

Moving from single loci to the descent of founder genome segments, it seems initially most natural to consider downward simulation, as for example in Brown et al. (2012). Generation by generation, parents can be selected to generate offspring, the pattern of meiosis across a genome realized, and the consequent labelling of segments by the appropriate haploid founder genome (FGL) recorded. Only the current generation of segments (with endpoints formed by past meiosis breakpoints), need be retained to generate each next generation. However, upwards simulation can be as or more efficient. First, every individual has exactly one father and one mother, but may have a quite variable number of offspring (Figure 1). Next, individuals with no current descendants need never be considered (Figure 1: right). More importantly, neither need any part of an ancestral genome that does not survive to the current generation. Since large parts of most ancestral genomes rapidly become extinct (Section 3.1), this is a very considerable saving.

**Figure 1:**
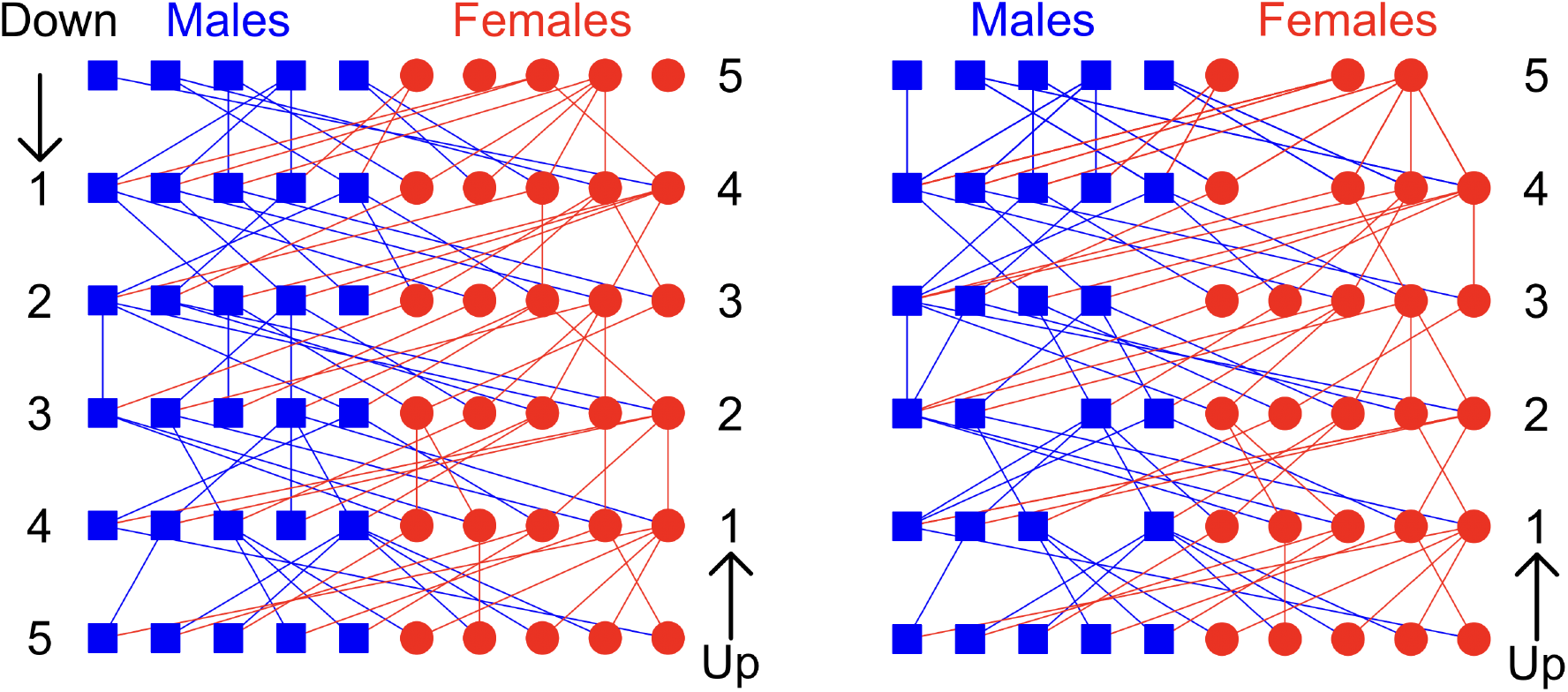
Genome ancestry and descent: up versus down

As when proceeding downwards, only the generation at *k* generations ago needs to be considered when proceeding up to generation (*k* + 1) (Figure 1: right). Pedigree information can be retained if, for example multiple realizations are required on the same pedigree, or if separate chromosomes are simulated separately on an given pedigree. In general, since all meioses are independent, meiosis breakpoints never need to be retained other than as their consequence as segment boundaries. The single exception is monozygous twins, who share both their maternal and their paternal meioses, with information from their descendants being processed jointly through these two meioses. On an arbitrary complex pedigree, ordering of individuals (or their genomes) is important. Consider again the grandfather-granddaughter mating, or even the mating of a (male) clone from an earlier generation to current individuals (Thompson, 2023). The maternal genomes of their offspring must be processed up to any relevant descendants of the male. The paternal genomes can only be processed once the paternal genomes of all offspring of the male are available. While a little more complex than downward generation of diploids by age order (parents preceding offspring), the same principle can be applied to the haploid genomes when simulating upwards.

Although upwards simulation is in many ways more practical, there are two slight disadvantages. The first is the bookkeeping of the set of descendants who will receive a precise segment of ancestral genome. As one proceeds to earlier generations there are a very large number of segments with changing sets of descendants determined by the meiosis breakpoints lower down the pedigree. Keeping count of the number of (unique) descendants of each segment is relatively straightforward (Section 2.4), but even then the number of segments increases very quickly. The second disadvantage is in comparing realizations over successive generations. Simulating downwards, the same founder FGL can be scored at 5, 10, 15, 20,,, generations on a single pedigree. Upwards, we are considering survival to the present from ancestors 5, 10, 15, 20 … generations ago. That is, they are different pedigrees, although with the same pedigree process. Note that when considering descent say from 10 generation ago, we will refer to the 10th generation ancestors as founders, and to their haploid genomes as FGL, even though they are an intermediate generation in a deeper pedigree of for example 20 generations. In this paper we will use both upwards and downwards simulation, as the occasion demands.

### 2.3 Ancestry of a single haploid genome

For an unbounded ancestral population and under a Haldane genetic map, Thompson (2024) described the processes to realize ancestry in a parental diploid genome, given the segments of ancestry in offspring chromosomes, and hence realize the ancestry of a current haploid genome – for example, an individual’s maternal genome.

Forwards in time, the meiosis process consists of parental chromosomes duplicating to form a tetrad, and crossovers then occurring, resulting in offspring chromosomes consisting of alternating segments of the parent’s maternal and paternal genome (Figure 2). In the reverse view, the offspring chromosome splits into segments in the parent’s maternal and paternal chromosomes. In tracing back the earlier history of these segments as shown in the lower part of Figure 2, there are two distinct processes to be considered. First ancestral segments may be broken by crossovers in the parental meiosis, as shown at the next earlier generation in Figure 2. Second, where there are multiple ancestral segments in a single chromosome, they may be separated by recombination (that is, an odd number of crossovers) in the *non-ancestral* intervening chromosome segment, as shown at the next earlier generation in Figure 2.

**Figure 2:**
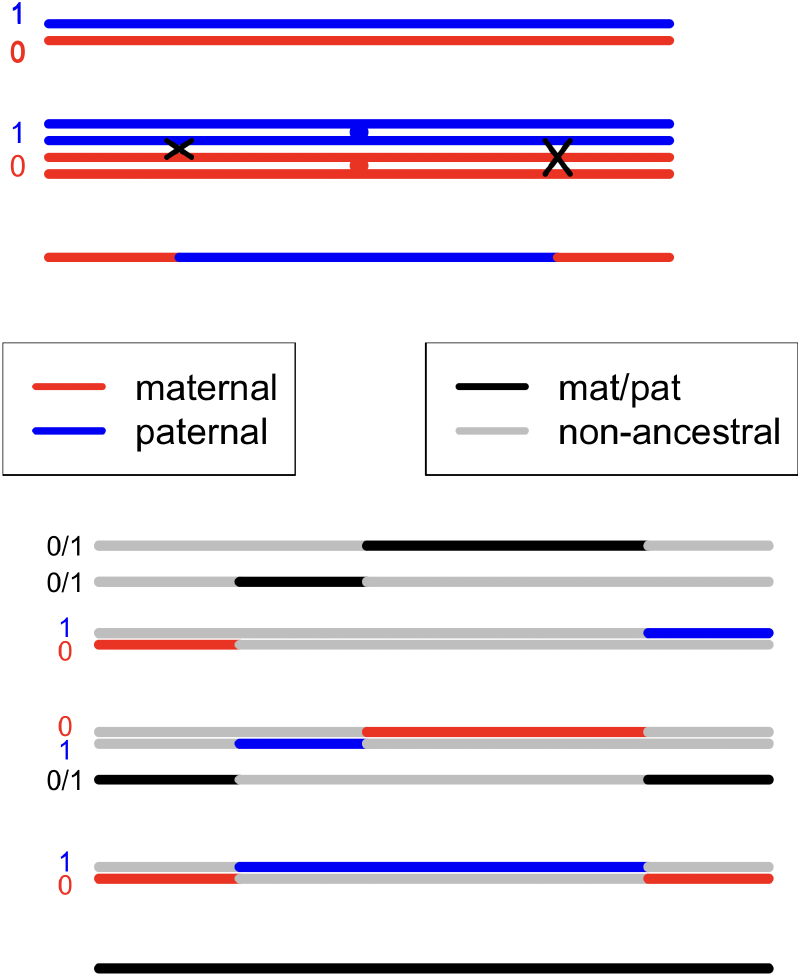
Above: crossovers occurring in a chromosome in meiosis. Below: the resulting processes of ancestry: see text for details. (Figure from Thompson (2024).)

For haploid genomes, define

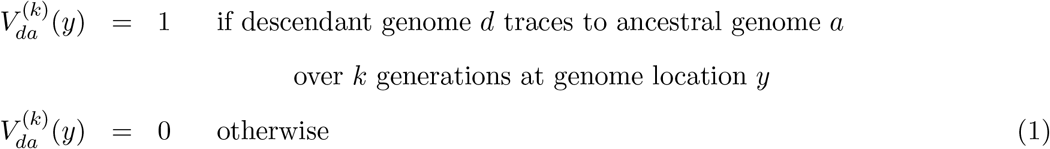

for *d* = 1, 2*, …,* 4*N*, *a* = 1, 2*, …*4*N*, *k* = 0, 1, 2, 3, 4*, …* and 0 *< y < L*. If we are considering upwards from from the current generation, 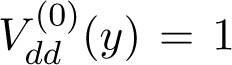 for each *d* and all *y*, and we count generations *k* = 0, 1, 2, 3*…* backwards in time. If we are considering downwards from a founder population, then 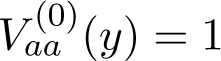 for each *a* and all *y*, and we count generations *k* = 0, 1, 2, 3*…* forwards in time. Note that for each point *y* and each generation *k*. for each descendant *d* there is a unique *a* for which 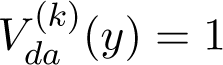. However, for given *a* there my be many *d* for which 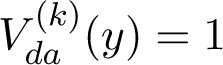. Note also that although we consider 0 *< y < L* as a continuum, the segmental nature of inheritance means that over 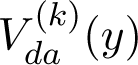 switches between 0 and 1 at a finite number of distinct points.

Although the focus of this paper is descent and survival of genome, we consider first the case of ancestry of a single haploid genome, since this will provide the method for considering descent through backward simulation of genome ancestry. The case of ancestry of a single genome was considered in Thompson (2024), but exact simulation was restricted to the case of an unbounded ancestral population.

Under a Haldane map function, the segments in an offspring may be ascended to a parent sequentially from the first to the last on that chromosome, in accordance with the genetic map, switch points points occurring as a Poisson process, both within and between segments. More generally, for a non-Haldane meiotic model, one can generate breakpoints across relevant portion of the chromosome in accordance with the desired model, and then ascend the segments according to the realized breakpoints.

In an unbounded ancestral population, the only case of multiple ancestral segments of a single haploid genome in a single chromosome is where recombination has broken a segment into more than two pieces, as in the example of Figure 2. Two segments within a chromosome share the same ancestral lineage. In a finite population, genetic ancestors may share parents, and the parental chromosomes may contain segments of different lineages (Thompson, 2024). However, this does not affect the realization of the ascent of segments from offspring to parental chromosomes, since the meioses from the parent to the offspring are independent. Nor can the segments overlap, since each point of the focal genome can be in only a single ancestor at each previous generation. (*I am indebted to Dr.Phil Green (Pers. comm.) for these insights.*) This provides for efficient exact simulation of segments of ancestry of a single haploid genome in a finite population.

An example is shown in Figure 3. On the left are three homologous maternal chromosomes of 3 distinct individuals at some past generation, with the black segments indicating ancestry of the focal descendant haploid genome. Also shown are realizations of the relevant meiotic events in the independent meioses from their mother. These are the maternal (0) or paternal (1) indicators of the meiosis at the start of the first ancestral segment, and crossover points (X) within the relevant portions of each chromosome. This information determines the segments of ancestry in the maternal (red) or paternal (blue) genome of each mother, shown in the central panel. Only then is it necessary to recognize shared parentage. If the three individuals share a single mother, then the segments of ancestry are simply combined into the maternal (red) and paternal (blue) chromosomes of the mother, as shown in the right panel of Figure 3. The segments in the mother will derive from different ancestral lineages, depending through which offspring they trace, but this does not impact their realization as ancestral segments of the focal genome, nor further ancestral tracing. Note it is irrelevant what the mother passes to each offspring in regions other than those shown in black: only those segments are ancestral to the focal descendant genome.

**Figure 3:**
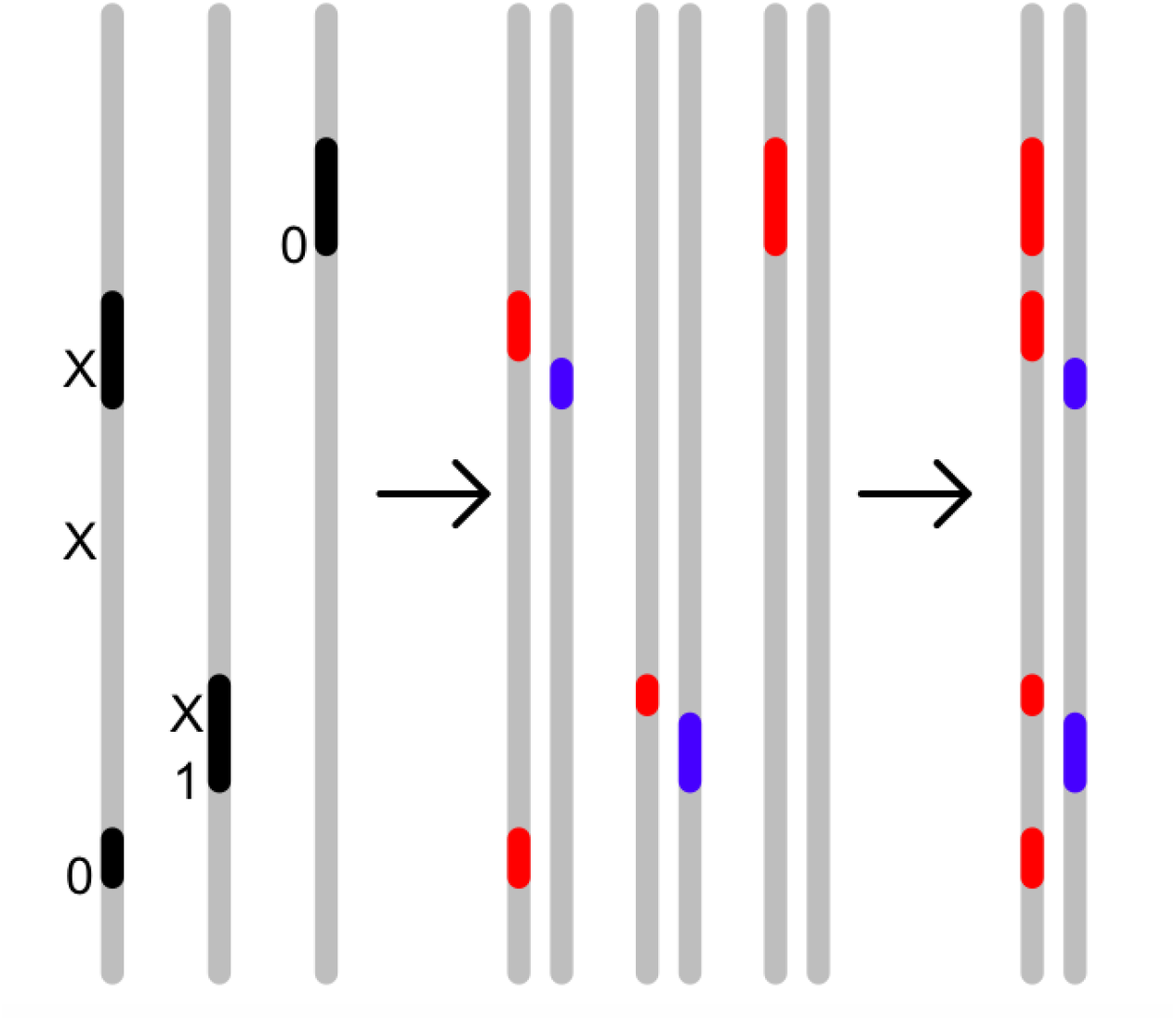
Realizing ancestry of segments in a 1 Morgan (100 cM) length of genome: for details see text

Given a set of ancestral segments in the maternal and paternal genomes of a set of ancestors, first one may generate the parents of those individuals under the population model. Given a region of a current chromosome with segments of ancestry, these may be ascended to the paternal and maternal chromosomes of the parent, with segments via different offspring being combined into the parent chromosomes as in Figure 3. Only parental chromosomes containing ancestral segments must continue to be traced: others can be dropped. Then, as for downward simulation, one simply proceeds to the next (earlier) generation.

There are a few caveats to this procedure. First monozygous twins are an exception to the procedure, since they share both their maternal and paternal meioses. Segments in monozygous twins must be ascended jointly to their parents’ chromosomes. The same restriction of course applies when considering downward descent to monozygous twins, but is more immediate in that case, since monozygous twins have identical ancestry. Second, in a pedigree without discrete generations and/or complex inbreeding, care must be taken that meioses are realized in ascending order, so that the relevant information is available at each step. Finally, the method applies only to the ancestry of a single haploid genome. To extend even to the ancestry of two current genomes, for example both the maternal and paternal genomes of a current individual, segments in ancestral chromosomes will be characterized by being ancestral to paternal, maternal, or both or neither. Rather than considering ascent of segments, it is simpler to generate the switch points in each meiosis, and retain them until both (or all) genomes have been ascended through the meiosis, as described in Section 2.1 above.

### 2.4 From ancestry to survival and descent

Using equation (1) we now define many quantities of interest to gene survival and descent of ancestral haploid genomes *a* to descendant haploid genomes *d*:

1. **Ancestry:** The length of genome in *a* that is ancestral to *d* at generation k is 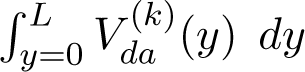. Genome *a* is a genetic ancestor of *d* if this quantity is greater than 0. Then number of genetic ancestors of *d* at generation *k*, is the count of *a* for which this quantity is greater than 0. Finally, the sum over *a* of this quantity is *L*, since genome *d* traces to a unique *a* at each 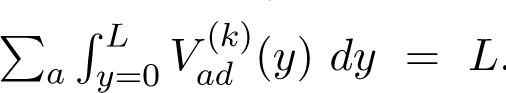.
2. **Survival:** Let 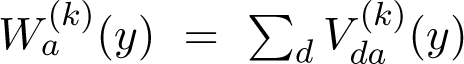 denote the number of copies of ancestral genome *a* at *y* at k generations later., and 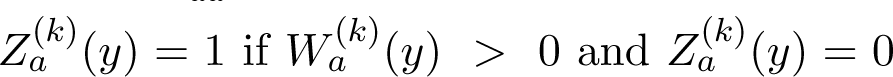 otherwise. Then 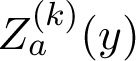 is the indicator function of survival of *a* at *y* from ancestral genomes *k* generations prior, and 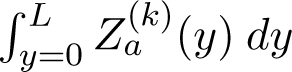 is the amount of the genome of *a* that is surviving.
3. **Descent:** The descent counts *W* ^(*k*)^_*a*_(*y*) may also be scored across the genome. For example, we may consider the total length of genome descent from *a*: 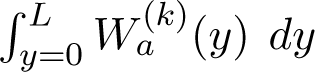 or the mean number of descendants at a location conditional on survival at that location:

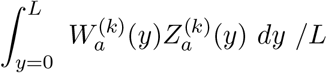

Alternatively, we may consider statistics such as the maximum number of descendants of *a*:

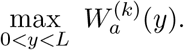

We provide the backward simulation details for survival 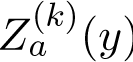 and then for descendant counts 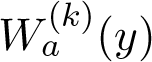 in a finite population using the same example as in Figure 3. In Figure 4, the genomes on the left again represent three maternal genomes at generation *k* and the black segments now represent genome surviving to any current (generation-0) genome. Of course, the segments now are larger, and may overlap: although each generation 0 point has a single ancestry at generation *k*, different generation 0 genomes may trace to different generation *k* ancestors.

**Figure 4:**
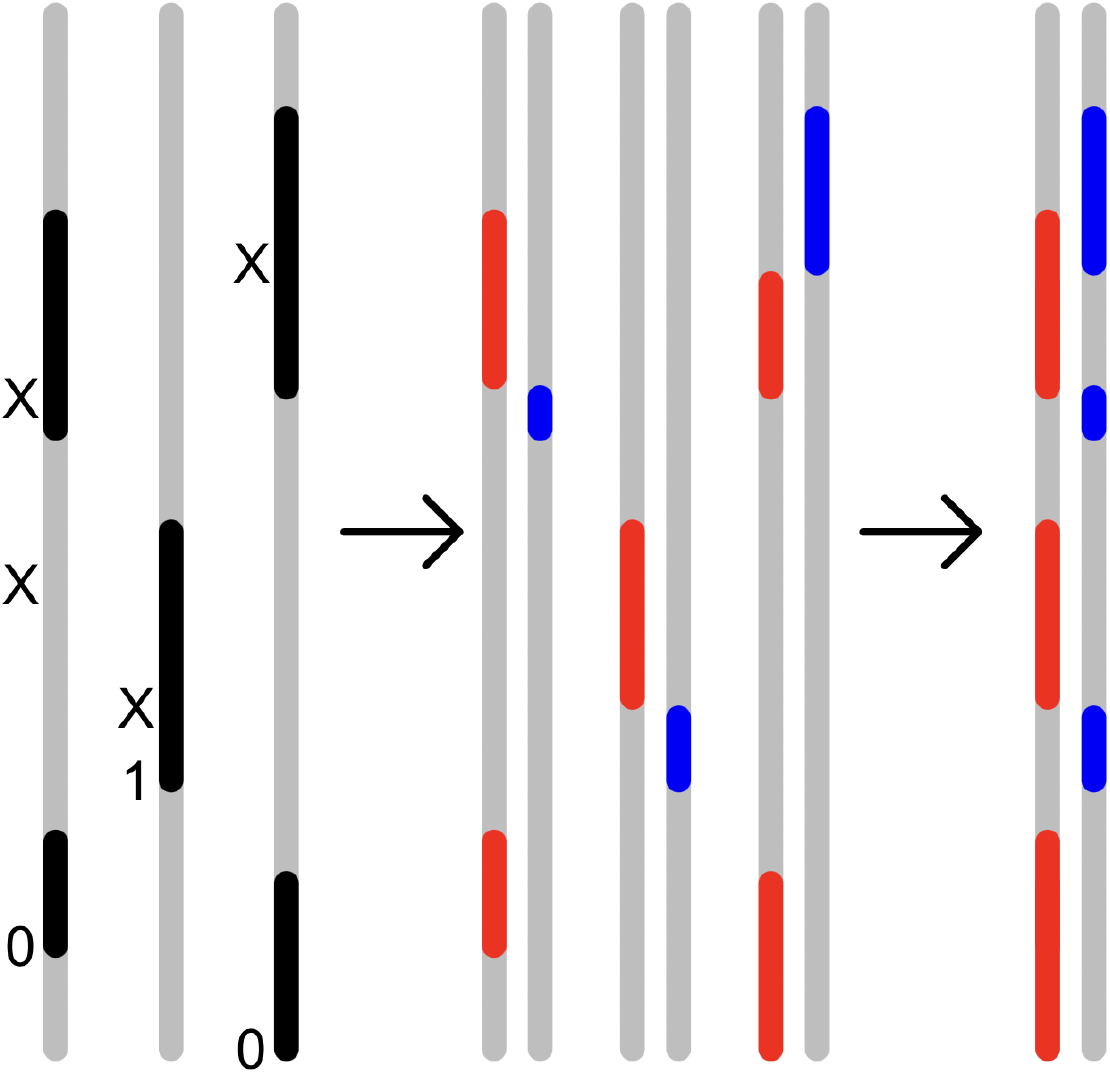
Realizing survival of ancestral genome in a 100 cM length of genome: for details see text

Genomes are ascended from the individuals to their mothers at generation (*k* + 1), just as before, and only then do we need to recognize if these three individuals have the same mother. The segments are then combined just as before: the mother’s paternal/maternal genome will survive at point *y* to generation 0, if it segregates to a surviving segment in any of her offspring.

A similar situation applies to descendant counts, but the situation is a little more complex. The surviving segments in the three generation-*k* individuals, will have subsegments with varying numbers of descendants due to meiosis breakpoints lower down the pedigree in generations 1 to (*k−*1) (Figure 5). However segments can be ascended separately to the three generation-(*k* + 1) maternal and paternal genomes. Now when it is found that these three mothers are the same generation (*k* + 1) individual, the counts can be combined, summing descendant counts in relevant subsections, since each generation 0 genome ascends at any location to precisely one generation *k* genome.

**Figure 5:**
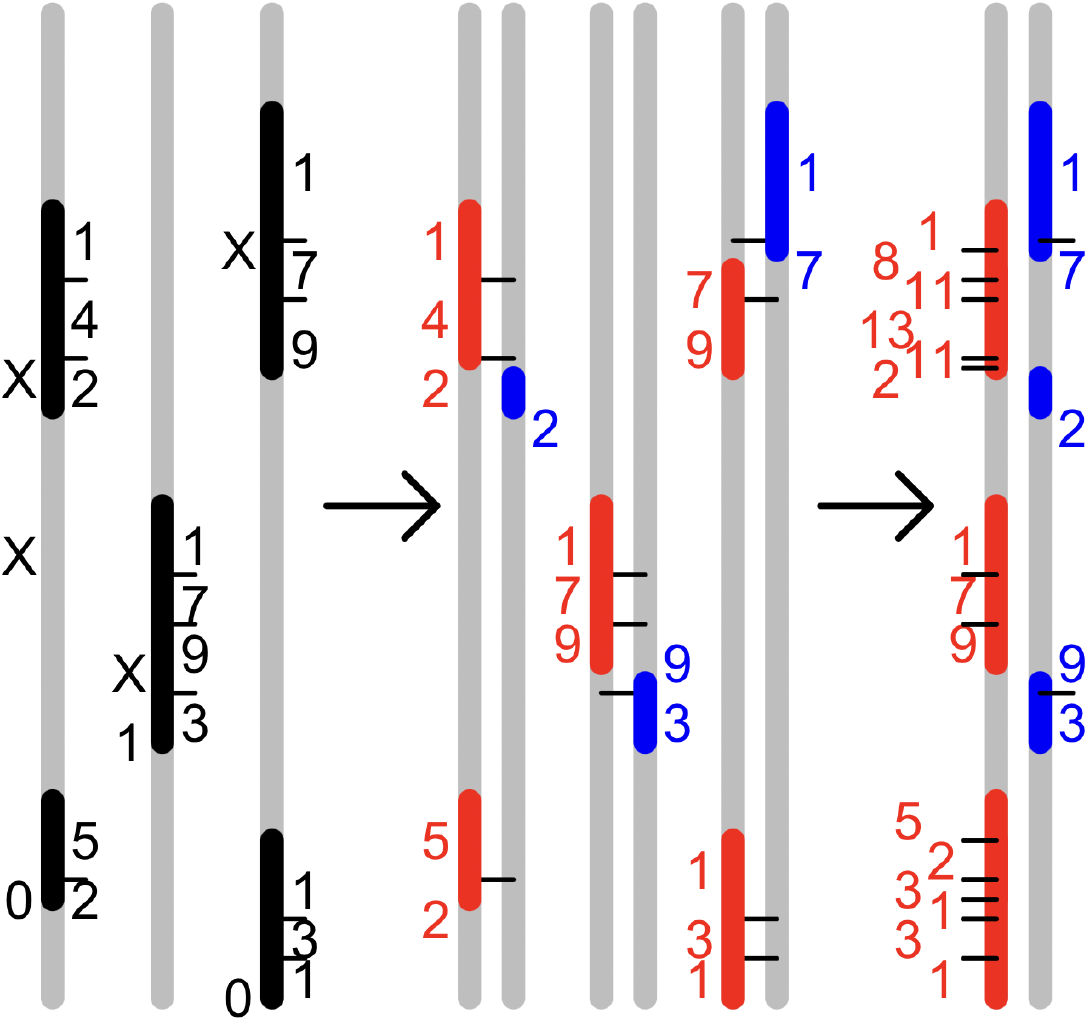
Realizing descendant counts of ancestral genome in a 100 cM length of genome: for details see text

For example, combining the counts (1,4,2) in the maternal genome of the mother of the first individual, with the overlapping counts (7,9) in the maternal genome of the mother of the third, we obtain a segment with 6 subsegments have counts (1, 8=1+7, 11=4+7, 13=4+9, 11=2+9, 2). [Note: the figure is slightly unclear: the kid1 red segment with a count of 2 extends from position (in cM) 63 to 65, while the kid3 red segment with a count of 9 extends from 64 to 71.]

### 2.5 Some basic results for unbounded populations

To provide a comparison with finite populations we consider first an unbounded populations. First we consider the count of ancestors and descendants, assuming every diploid individual has independently a Poisson (mean 2) number of offspring. Each individual has 2*^k^* ancestors, *k* generations ago: there is no variation. For descent, the expected number of descendants *X_k_*at *k* generations is also 2*^k^* but there is substantial variation:

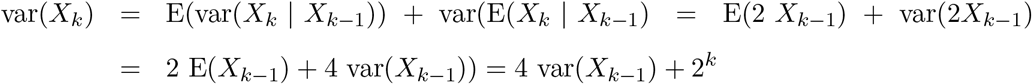

Solving with var(*X*_1_) = E(*X*_1_) = 2, we obtain var(*X_k_*) = 2*^k^*(2*^k^ −* 1). In a finite population of course both numbers of ancestors and of descendants are limited, but we will see this same pattern of difference between ancestry and descent. This of course relates again to the fact that every individuals has one father and one mother, but a variable number of offspring.

Next we consider extinction, using standard branching process theory. If the probability generating function of the offspring number *N* at generation *k* into the future is *h_k_*(*z*) = E(*zN*) then the probability of extinction *q_k_* of the branching process by generation *k* is given iteratively by

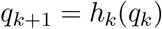

For diploid individuals, the offspring distribution is Poisson with mean 2, so *h_k_*(*z*) = exp(2(*z −* 1)) for all *k*, and *q_k_* converges rapidly to 0.203. The long-term probability of descendants is just about 80%, and the existence of such descendants is effectively determined within about 5 generations. For a single haploid gene, with 50% chance of passage at each meiosis, the offspring distribution is Poison with mean 1, so *h_k_*(*z*) = exp(*z −* 1). The extinction probability converges to 1. However convergence is slow, reaching 91.2% at 20 generations and 95.4% at 40 generations.

For the number of copies of a single genome location, the number of copies at the next generation is Poisson (mean1), and the expected number of copies at each subsequent generation is 1. Derived similarly to above, the variance after *k* generations is *k*. Note that in any population, finite or unbounded, and under any pedigree model, the expected proportion of any length of a founder haploid genome that survives is simply the point-wise probability of survival. If the expected descendant count is 1, then the expected number of descendant copies conditional on survival is the inverse of the probability of survival.

The extinction probabilities of diploid individual descendants (red) and haploid gene copies (black) are shown on the left side of Figure 6. The results for genomes of varying length are of course bounded by these two curves. By simulation of the counts of subsegments of genome in the genomes of each diploid descendant, extinction of lengths 10cM, 1M, 2M, 2 lengths of 1M, 10M, and 30M are also shown in Figure 6. On the left is shown the extinction probability over up to 40 generations. The right-hand figure shows the same information for the same 7 genome lengths (plotted on the x-axis on a log_10_ scale), at generations 5, 20, 40 and 100.

**Figure 6:**
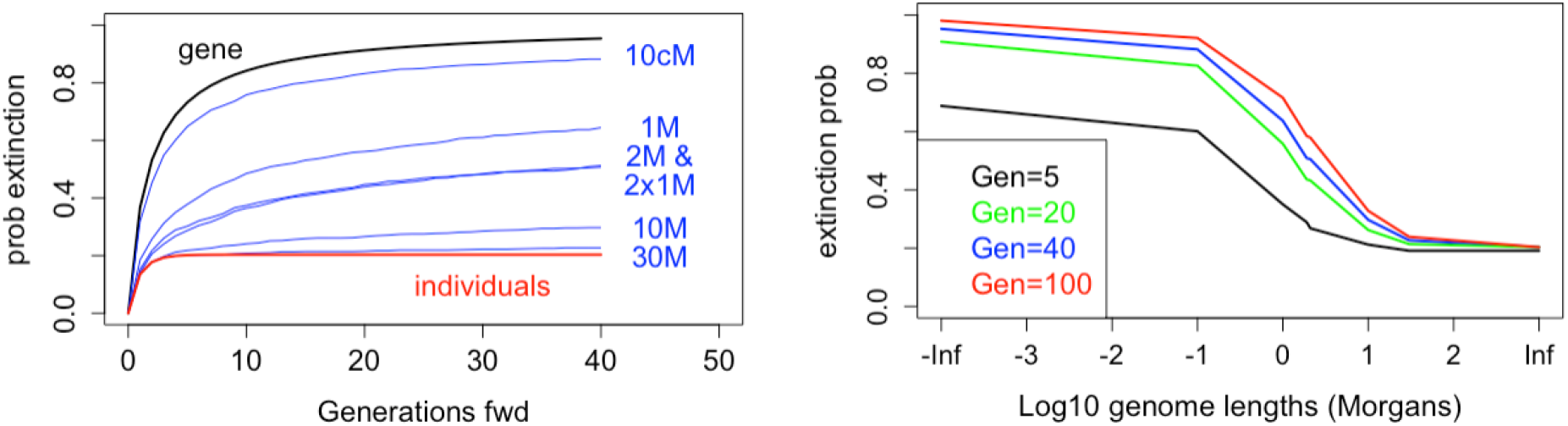
Probabilities of extinction: Unbounded population

A segment of length 10cM (0.1M: of order 10^7^bp) has lower extinction probability than a single point, but because of the segmental inheritance of DNA, still has high probability of extinction. In the first few generations it behaves almost like a single point in the genome. Only as it gets broken up and dispersed does the probability of survival of some part of it diverge from the single point curve. At the other extreme, for a 30 Morgan genome the probability that some part will survive is close to the probability of having descendants over 40 generations. However it is possible to have descendants but no genetic descendants, and the extinction probability for a 30M genome continues to increase over time.

A genome length of 1 to 2 Morgans (typical of a mammalian chromosome) has an extinction probability around 50% over 40 generations, but these extinction probabilities also continue to increase over time. Also shown in Figure 6 are two lines, one for a segment length 2 Morgans, and the other for two 1-Morgan lengths. As expected, the latter has a slightly lower extinction probability in the early generations, but overall the exact division of genome into chromosomes has negligible effect. As with many other results we see the major change between 5 generations and 20 generations (Thompson, 2024). There is relatively little change between 20 and 100 generations, but note that extinction of any finite length of genome must continue to rise, since the expected proportion of genome surviving continues to decrease.

## 3 Survival and descent of individuals, genes, and genomes

Throughout this section we will consider a finite random-mating population. Numerical results will be presented for dipolid populations of 100 males and 100 females in each generation. We use the classic model of no genetic interference for meiosis (Haldane, 1919). Issues of population size, mating structure, and family size distributions will be considered in Section 4.

### 3.1 Extinction, ancestry, and descent of individuals in a finite population

#### 3.1.1 Extinction of haploids and diploids

For a single genome location (0-length genome segment) or for individuals, the generation to generation probabilities of descendant count (and hence of extinction) are straightforward. In a random-mating population of 2*N* individuals (4*N* genes), suppose at generation *k* (forwards in time) a single diploid founder (generation 0) individual has *F_k_* female and *M_k_* male descendants, and *D_k_*= *M_k_* + *F_k_*. The probability that a diploid individual at the next generation has a mother that is not a descendant of the focal founder is (1 *− M_k_/N*), and similarly for fathers, with all individuals being independently assigned to parents. Thus the probability that an individual is not a descendant of the focal founder is

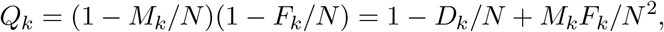

and the number of descendants at the next generation is *Bin*(2*N,* (1 *− Q_k_*)). The number of females *F_k_*_+1_ among these descendants is a hypergeometric sample from the total *N* females and *N* males from the next generation, and *M_k_*_+1_ = 2*N − F_k_*_+1_.

For the haploid genomes of founders (FGL), suppose that generation *k* (forwards in time) at a given genome location, an FGL has *d_k_* descendant copies; *f_k_* in females and *m_k_* in males. Again, offspring are assigned independently to parents, so the probability each next-generation maternal [paternal] gene descends from this FGL is *f_k_/*2*N* [*m_k_/*2*N*] regardless of how the parent copies are distributed among the *N* female [male] parents. The number of copies in the offspring distribution is the sum of two independent binomials.

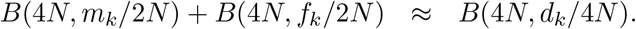

Strictly the two binomials have slightly different probability parameters, so the total is not Binomial. However, at each generation the population is constrained to have equal numbers of males and females in total, and parents are randomly assigned. For practical purposes the approximation *B*(4*N, d_k_/*4*N*) suffices.

The process is easily simulated. Using 1,000,000 realizations for *N* = 100, Figure 7 shows the simulation values for the finite population superimposed as black dashed lines on the theoretical blue and red extinction probabilities for an infinite population (see the outer lines of Figure 6 (left panel)). It is clear that differences in extinction between the infinite and this finite population are undetectable on this time-scale: see also Section 4.1.1. As in Section 2.5 note that the point-wise probability of FGL extinction is also the expected proportion of genome that is extinct.

**Figure 7:**
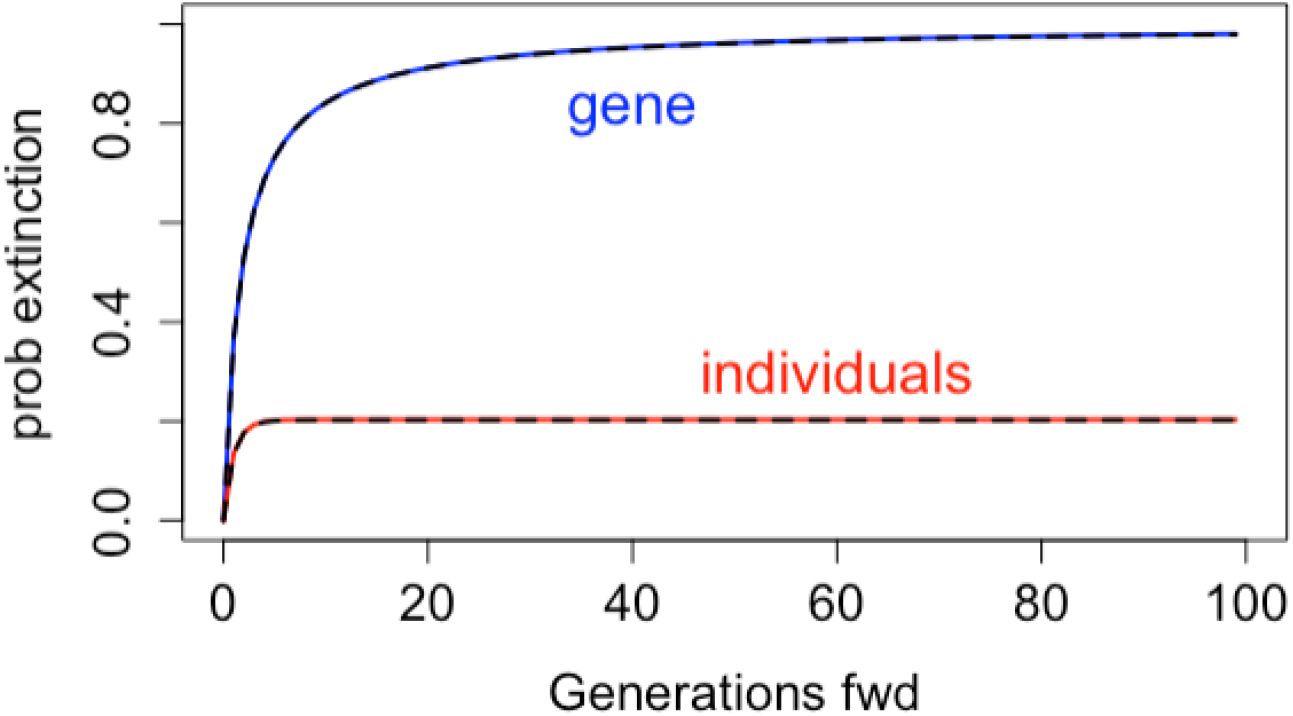
Probabilities of extinction in a small population. Comparison of the infinite population (red and blue lines), with a random-mating population of 100 males and 100 females (dashed black lines).

#### 3.1.2 Individual diploid ancestry in a small population

Genome segments of ancestry or descent exist within the genomes of the diploid individuals of the population. Thus first we consider the processes of diploid ancestry and descent in a finite random mating population. Consider a population of 2*N* diploid individuals with *N* males and *N* females at each generation. We first consider the marginal distribution at *k* generations ago of the number of ancestors *A_k_* of a single current individual.

Note that *A*_1_ = 2 and that in expectation *A_k_* increases to approximately 80% of the total population; E(*A_t_*) *→* 1.6 *N* . In the left panel of Figure 8 is shown the result of 200 realizations of the process for a population of size 200, with the blue curve showing the empirical mean. Within about 12 generations, the mean is very close to 160.

**Figure 8:**
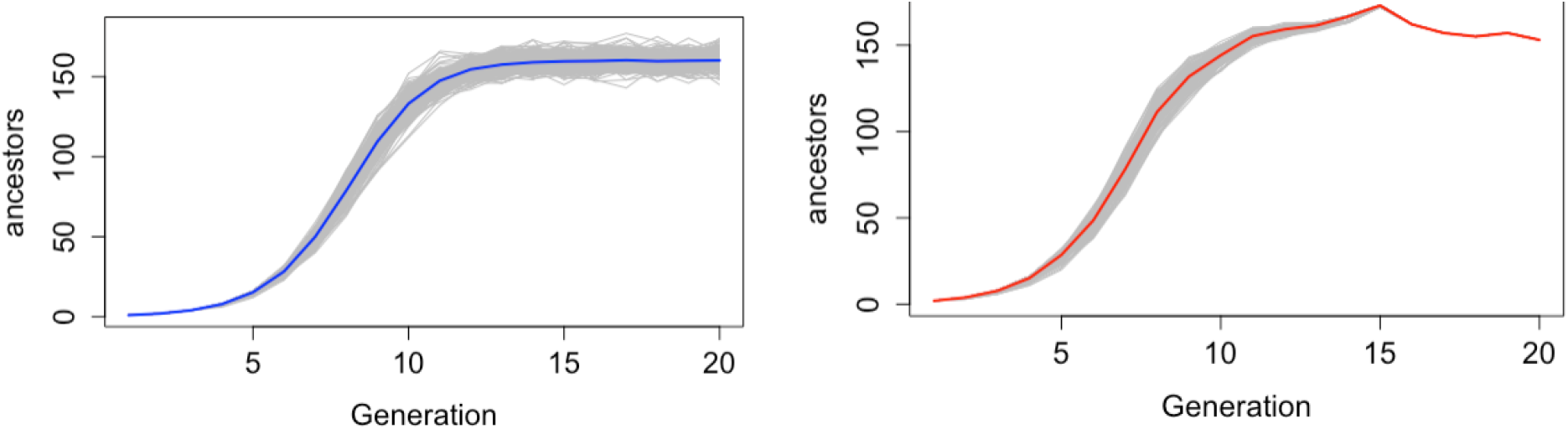
Count of ancestors a random mating populations of 100 males and 100 females:, with generations counted backwards in time: (Left) 200 independent realizations;: Right: : 200 individuals of a single pedigree realization.

In contrast the right panel of Figure 8 shows the counts of ancestors for the 200 members of a single population pedigree generated by random mating of males and females. Within the population, within 17 generations, all individuals have exactly the same ancestors, and thereafter the count varies about the expected 160 value. Even before reaching this limit, after about 10 generations the within-pedigree variation is much smaller than the total variance, there is strong dependence among individuals in the identity of their ancestors. Figure 9 shows the empirical standard deviations, for the two sets of realizations in Figure 8.

**Figure 9:**
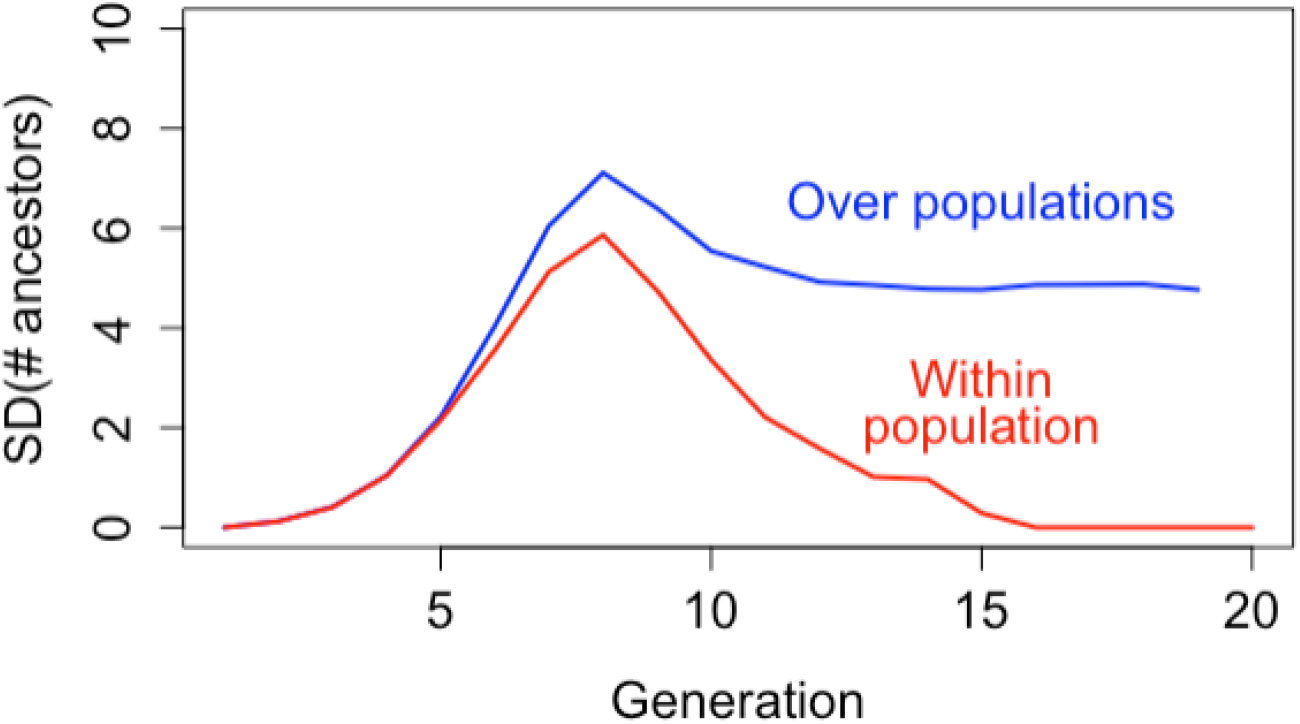
Empirical standard deviations in the count of ancestors.

Note that any exact formulae are complicated, and not very useful. If an individual has *A* ancestors at generation *k*, then these can be randomly assigned to the *N* females of the preceding generation, and the number of such females *F* who **not** ancestors (i.e. not mothers of any of the *A* individuals) is then given by the solution to the number of empty cells in the classical occupancy problem (Feller (1950), Section 4.5) assigning *A* items to *N* cells. Using the inclusion-exclusion probability formula for the union of events the result reduces to

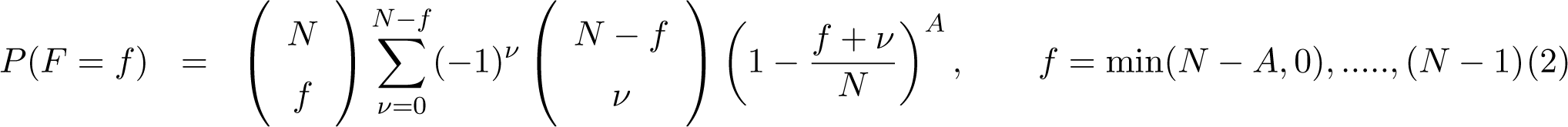

A similar formula independently applies to the number of males *M* at the preceding generation who are **not** fathers of any of the *A* ancestors, and the number of ancestors of the individual at the preceding generations is 2*N − F − M* . An alternative approach to this result is given in Supplementary Section A.

#### 3.1.3 Individual diploid descent in a small population

Descent shows a very different picture to ancestry, with much higher variance. The left panel of Figure 10 shows the number of descendants at each generation for 200 independent realizations of the process in a random mating population of 100 males and 100 females. Notes that a few individuals achieve ancestry of the entire population within 6 or 7 generations, while the approximately 20% of lines that become extinct do so within that same time frame., However, there are a few instances where the founder has few descendants even at 10 generation, and takes until 16 or 17 generations to become ancestral to the entire population.

**Figure 10:**
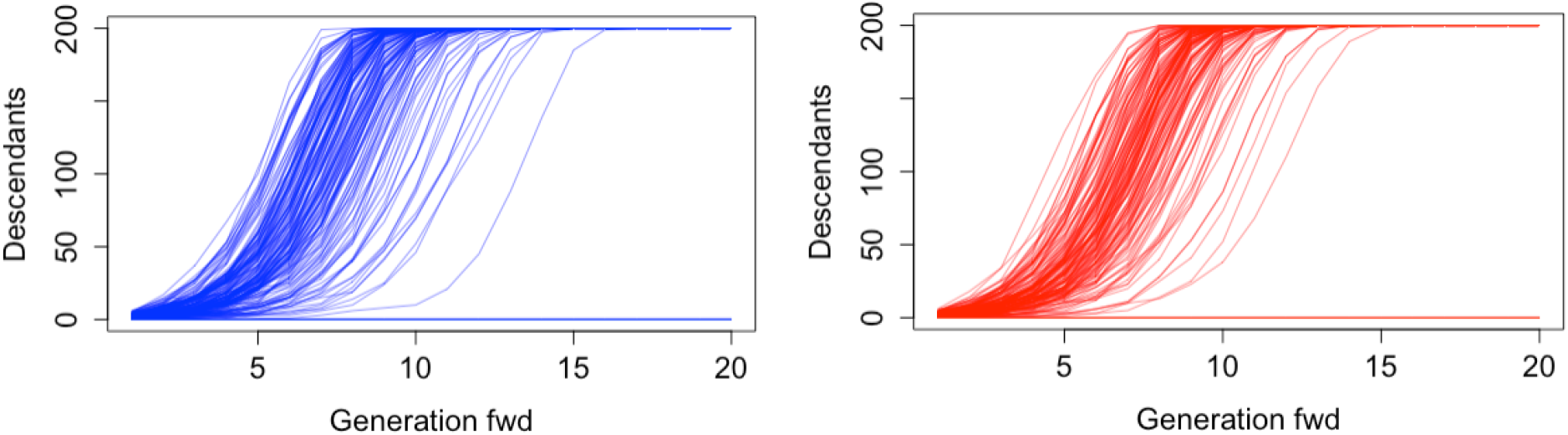
Count of descendants of an individual in a random mating population of 100 males and 100 females with generations measured forwards in time.: Left 200 independent realizations. Right The 200 members in a specific population pedigree.

As with extinction (Section 3.1.1) we may consider the the generation to generation process.. Suppose at generation *k* (forwards in time) a single founder (generation 0) individual has *F_k_*female and *M_k_*male descendants. The probability that an individual at the next generation has a mother that is not a descendant of the focal founder is simply (1 *− M_k_/N*), and similarly of fathers, with all parents being independently assigned to parents. Thus the probability that an individual is not a descendant of the focal founder is *Q_k_* = (1 *− M_k_/N*)(1 *− F_k_/N*), and each of *M_k_*_+1_ and *F_k_*_+1_ is binomial *Bin*(*N,* (1 *− Q_k_*). The process is thus easy to simulate, and the generation to generation means and variances are straightforward:

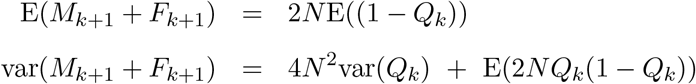

however explicit cumulative formulae are not feasible.

The right panel of Figure 10 shows the counts of descendants for the 200 founders of the same pedigree as shown for ancestry in the right panel of Figure 8. The two sets of curves in Figure 10 are similar, although the within-population curves have less variance, indicating there is some within-population dependence. Overall, the among-individual variance in descendant count is dominated by the approximately 20% of individuals who within about 5 generations have no descendants, so in Figure 11 we show the empirical standard deviations among individuals in their counts of descendant, conditional on them having surviving descendants. (The population curve (blue) is computed from a large number of realizations, not the specific curves of Figure 10.)

**Figure 11:**
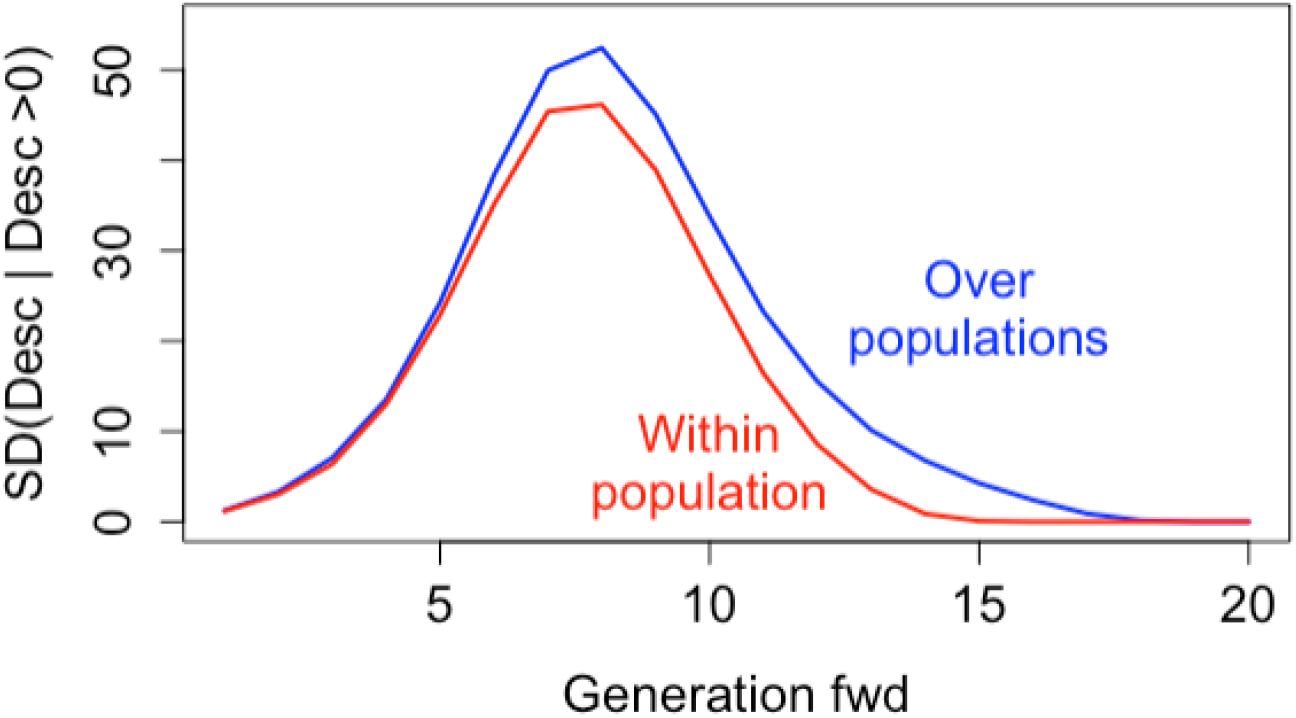
Empirical standard deviations in the count of descendants among founder individuals who have surviving descendants

In Figure 11 we see that for the first 6 or 7 generations there is almost no difference between the within- and between-population cases, and overall the within-population standard deviation is not much smaller than the between-population standard deviation. Both tend rapidly to 0 after about 15 to 17 generations: at this point all founders with surviving descendants are ancestors of the entire population, However, within a population, founders are a bit more similar as to the time-point at which this universal ancestry is achieved. In this example, the within-population SD becomes zero at 14 generations, while it takes 17 generations for the same number of independent realizations of descent to reach this point Comparing figures 11 and 9 we see both the greater within-pedigree dependence of ancestor counts compared to descendant counts, and also the much greater variance of descendant count overall. Due to the variance in offspring number, some individuals will rapidly have large numbers of descendants.

### 3.2 The descent of founder genome

In this section we will present results for survival and descent of genome in a single population pedigree of 100 males, and 100 females, generated by random mating. We consider a haploid genome of length 30 Morgans, divided into 30 chromosomes each of length 1 Morgan. Although only a single pedigree is used, we have already seen that variation over the individuals of a population of this size differs little from variation over the population (Section 3.1.3) and we will see in Section 3.3.2 that measures of variation within a population pedigree is little different from variation over pedigrees. Of course every population pedigree will have its extreme trajectories, and the particular instance will differ among pedigrees. Various particular extremes for our population are described in Supplementary Section B, as examples of what may occur, rather than as results that will occur in every population of this size.

#### 3.2.1 Segments of founder genome

In considering the decent of genome, the junctions of FGL formed by meiosis breakpoints thereafter descend as single point locations, and most become rapidly extinct. However, considering the same process by ancestry, every point in each current (haploid) genome traces uniquely to a genome in each earlier generation. Thus all the meiosis breakpoints retain an ancestry as boundary points of a segment for some FGL in some current individual(s). Hence for 2*N* individuals (4*N* founder genomes), each of length *L* Morgans, subdivided into *C* chromosomes, after *k* generations the total number of segments of founder genome in the population is Poisson Po(4*N* (*C* + *kL*)), or Po(4*N* ((*k* + 1)*L*) if *C* = *L*. To a remarkably close approximation each has an exponential length, mean 1*/*(*k* + 1) (Thompson, 2024), and the overall probability a segment is broken at the next generation is

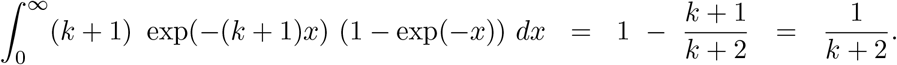

We consider now the way in which the Po(4*N* (*k* +1)*L*) total descendants genome segments at generation *k* are attributable to each of the 2*N* = 400 haploid founder genomes (FGL), and how many of the potential 2*N* = 400 current genomes each of the 400 FGL is represented in. In the population pedigree, a 30M genome was simulated downwards, using the *ibd create* program of MORGAN Ver 3.4 (2017) and the population genomes are output at 10-generation intervals from 10 to 100. Figure 12 shows the founder segment counts in increasing order at each of these output generations. First, just over 20% of founder genomes rapidly become extinct. This extinction probability is very similar to that is an infinite population. They have no current segments (left) and are present in no current genomes (right). Initially (at 10 generations) the number of descendant segments is fairly uniform among founder genomes with genetic descendants (left) as is the count of the genetic descendant genomes (right). Over time, there arise a few genomes with a very large number of descendant segments. At 10 generations the maximum segment count for any FGL is about 1300, but by 20 generations it is 3500, and by 100 generations there are FGL with almost 14,000 descendant segments, although the median, remains close to 2000. Of course, many of these are segments descending from the same portions of the founder genome. It is usually not the case that a large portion of the founder genome survives: see Section 3.3.1. Even at 100 generations some founder genomes have few surviving descendant segments and there are non-extinct founder genomes with segments in only a few current individuals. However, by this time,, 50% of all FGL (or 63% of surviving FGL) are present in all 400 current genomes.. At all generations, FGL with more than about 1200 segments are typically represented in all 400 current genomes.

**Figure 12:**
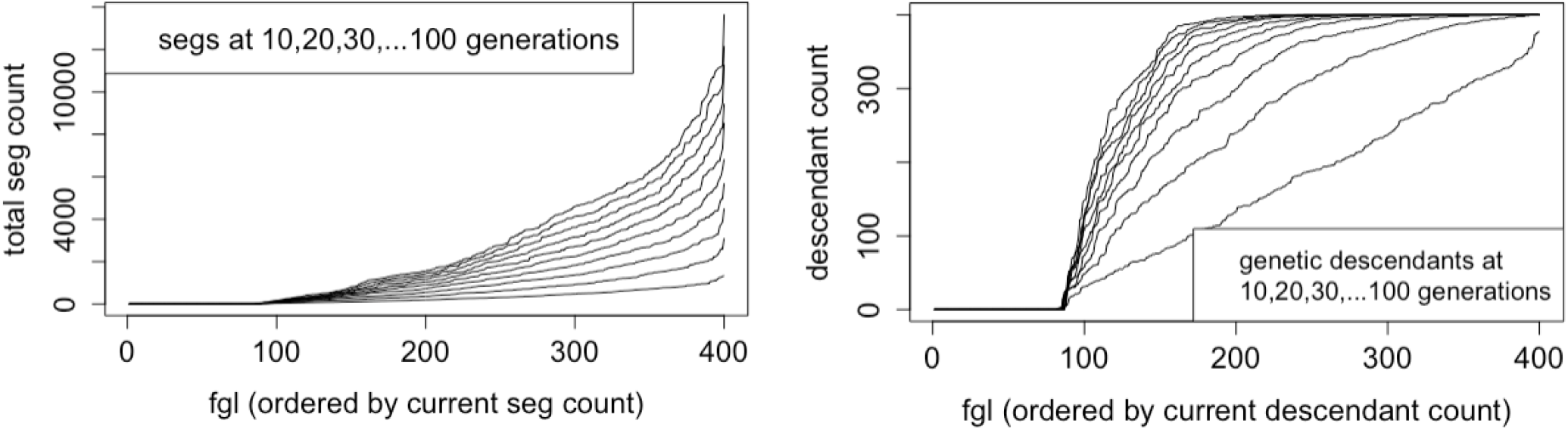
Descendant counts of the 400 founder genomes at intervals of 10 generations. Left: Ordered counts of descendant segments. Right: Ordered counts of current genomes that each founder genome is represented in.

Figure 13 gives a slightly different view of the same results, with descendants scored by diploid individual rather than by haploid genome. For a single random-mating population of 100 males and 100 females, in red are shown the number of descendants over the generations. (These are analogous to the red curves in Figure 10 (right) extended now to 40 generations.) In blue are the numbers of genetic descendants –diploid individuals carrying DNA from each founder.

**Figure 13:**
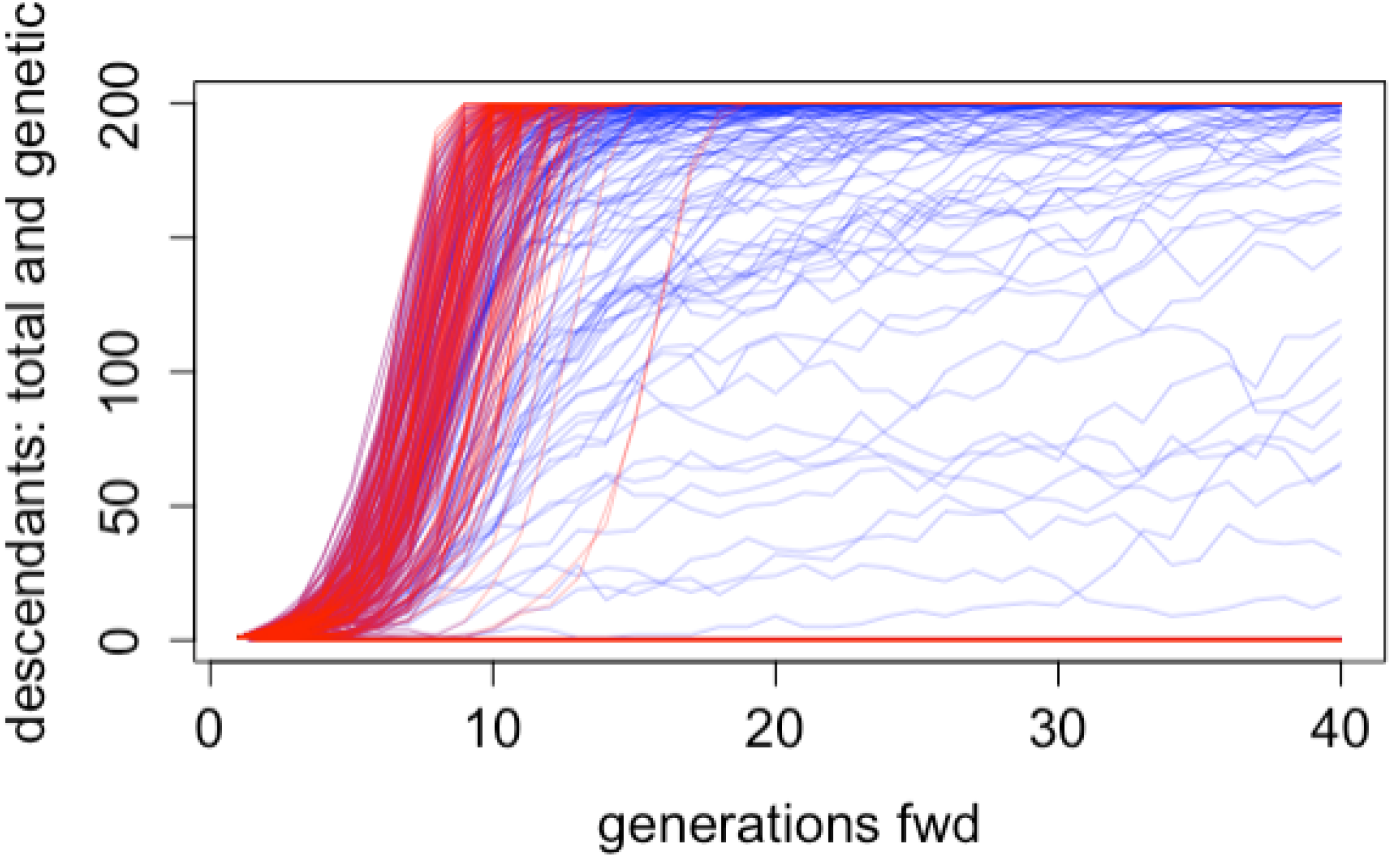
Comparison of the number of descendants vs number of genetic descendants.

Counts of genetic descendants rise more slowly than counts of descendants. Although many founders are genetic ancestors of almost all 200 individuals by generation 15, there are about 10 individuals who are ancestors of everyone, but are genetic ancestors of few. At the other extreme despite the existence of FGL with almost 14,000 descendant segments, distributed over a surviving genome length of only a few Morgans, no FGL are fixed at 100 generations, with very few FGL reaching a frequency of 75%. (Supplementary Figure B.3). More discussion of individuals with extreme low or high descendant counts is given in Supplementary Section B.

#### 3.2.2 Lengths of surviving and descendant genome

While descendant segment counts give a useful picture of survival of genome, it is possible to also score the Morgan lengths of surviving and descendant genome. In Figure 14 are shown the ordered values for each of the 30-Morgan 400 FGL surviving at 10, 20, 30, and 40 generations. On the left is shown the length of surviving genome; that is, the amount of each 30M founder genome surviving anywhere in the population. Surviving genome must necessarily decrease over time: the most successful FGL retaining about 55%, 33%, 25% and 15% of their genome surviving at 10. 20, 30 and 40 generations. As before 20% of FGL are extinct, and for those surviving in smallest amounts there is little change in these amounts after about 20 generations. On the right is shown the descendant genome, that is the total length of genome in the population that descends from that FGL at these generations. This shows a reversal at the extremes of largest genome descent, with the largest descent amounts increasing over time. There are a few FGL that have segments achieving polymorphic frequencies, and these have slightly increasing total descent over time; the maximal value is 125 Morgans at generation 10, 135 Morgans by generation 20, and 155 Morgans by generation 40. However note that only these extremes show change over time: most of the ordered values remain approximately constant. Although some founder genome continues to go extinct the same potential total 12,000 Morgans in the 400 genomes of founders are distributed among the total 12,000 Morgans in each generation. The extremes of these distributions are again discussed in Supplementary Section B.

**Figure 14:**
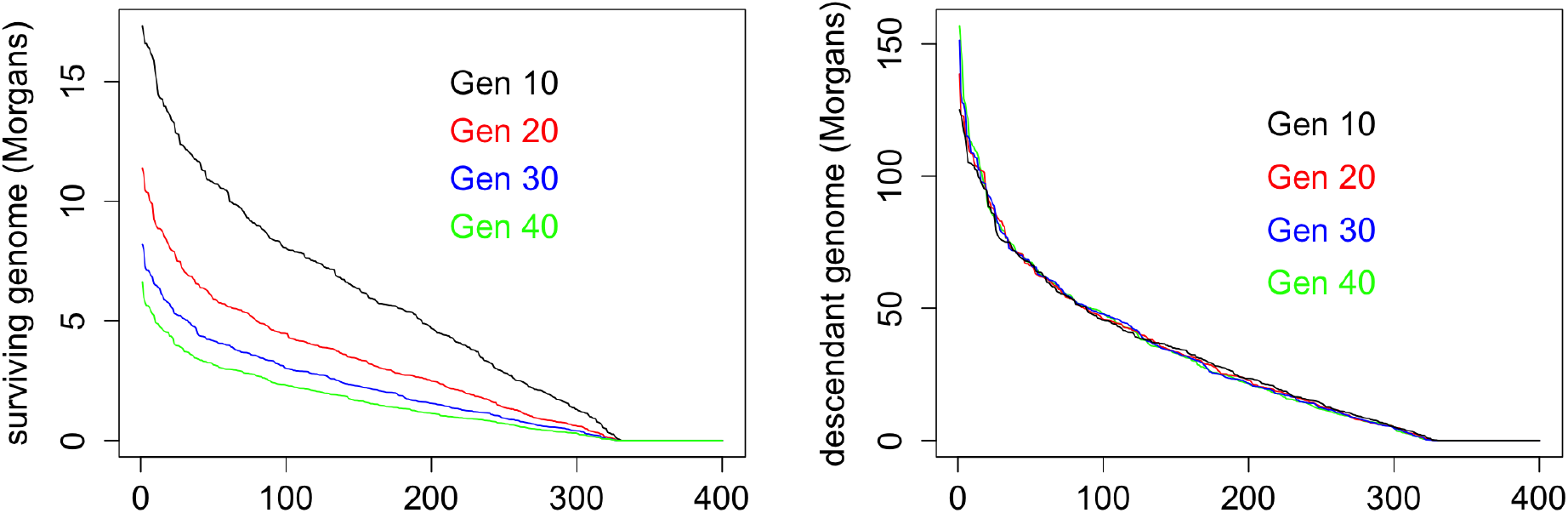
Surviving and descendant lengths of genome: for details see text

Figure 15 explores the relationship between surviving and descendant genome over time. On the left are shown the raw values for each of the 400 FGL. The correlation between surviving genome and descendant genome is high, rising from 0.91 at generation 3, to 0.95 by generation 7, and 0.98 by generation 40, although there is some variation. The ratio of descendant to surviving genome is a measure of the extent to which each founder genome is (on average) replicated in the descendant population. On the right this ratio is plotted over the 40 generations of descent from the founders. The FGL are colored according to their descendant genome at 40 generations, from the minimum (cyan) to the maximum (red) – the bar on the figure shows the range from minimum to maximum. The order of plotting the lines on the page is important. To minimize obscuring the results by over-plotting, lines are plotted from the central color to the extremes: that is in order (200,201,199,202,198,203, …, 2,399,1,400). There is still positive correlation between the ratio and total descent, but it is not nearly so strong as the raw correlations. The correlation of the ratio with descendant genome now decreases from about 0.9 at generation 5, to 0.6 by generations 20, and 0.55 by generation 40. Generally FGL with low replicate counts have low total genome descent, but there are a few genomes where the total amount of descendant genome and of surviving genome is low, but that genome is replicated in many current descendant genomes. This is further discussed in Supplementary Section B.

**Figure 15:**
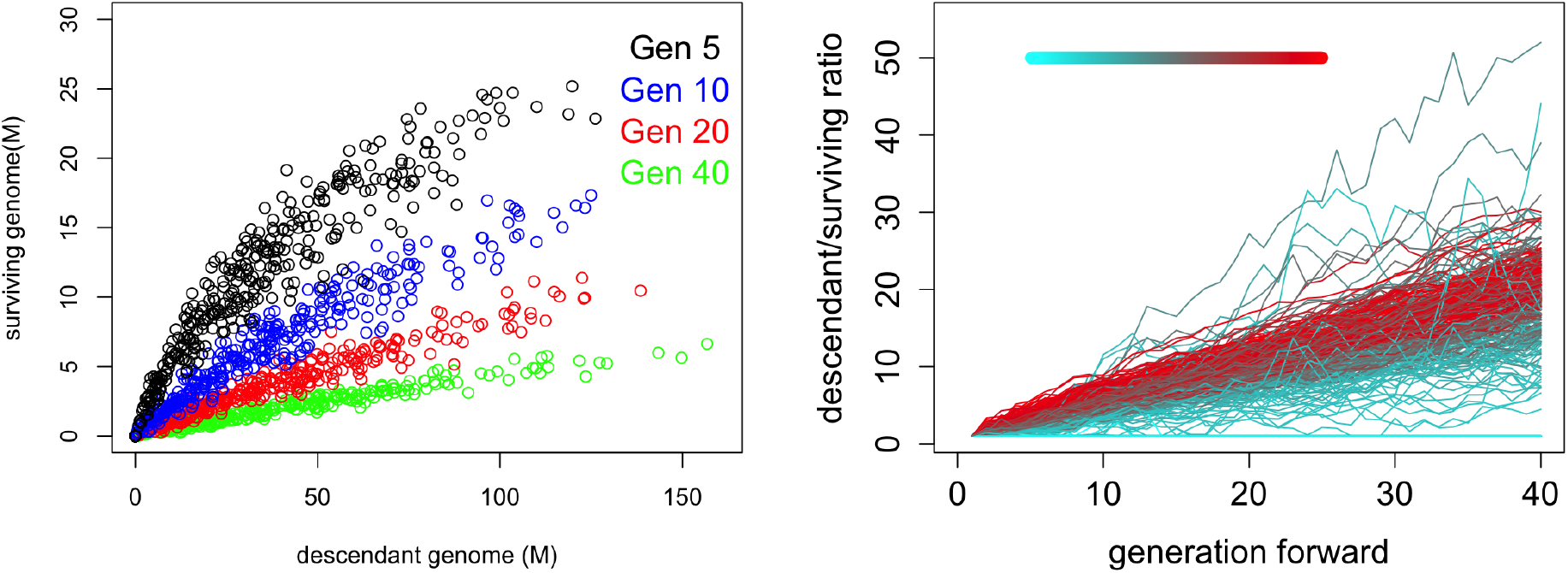
Comparison of descendant to surviving genome in individuals over time: for details see text

### 3.3 Descent of a 1 Morgan length of genome

In this section we provide two examples using the ancestry simulation methods of Section 2.4, with a focus on variation among founders, both within a given population pedigree and among populations. We focus on a length of genome of 1 Morgan (100 cM), for example a single chromosome. Again we consider the pedigree of a random mating population of 100 males and 100 females in each generation.

#### 3.3.1 Survival of genome

Consider first the length of each founder’s genome surviving over the generations. Due to the high positive dependence in survival of the genomes within a diploid founder (Section 3.4) we score the amount of genome surviving from each founder diploid (200 cM), although of course simulation proceeds via each individual’s haploid maternal and paternal genome.

Figure 16 shows on the left the distribution of the mean surviving genome length for each of the 200 diploid founders from 500 realizations on a given population pedigree, in decreasing order of those mean values. The results confirm that 20% of individuals become rapidly extinct, and the increasing loss of genome over the generations. Among the 160 diploid founders with descendants the median surviving genome decreases rapidly to about 65 cM by generation 5, 40 cM by generation 10, and thereafter more slowly to about 25 cM by generation 20.

**Figure 16:**
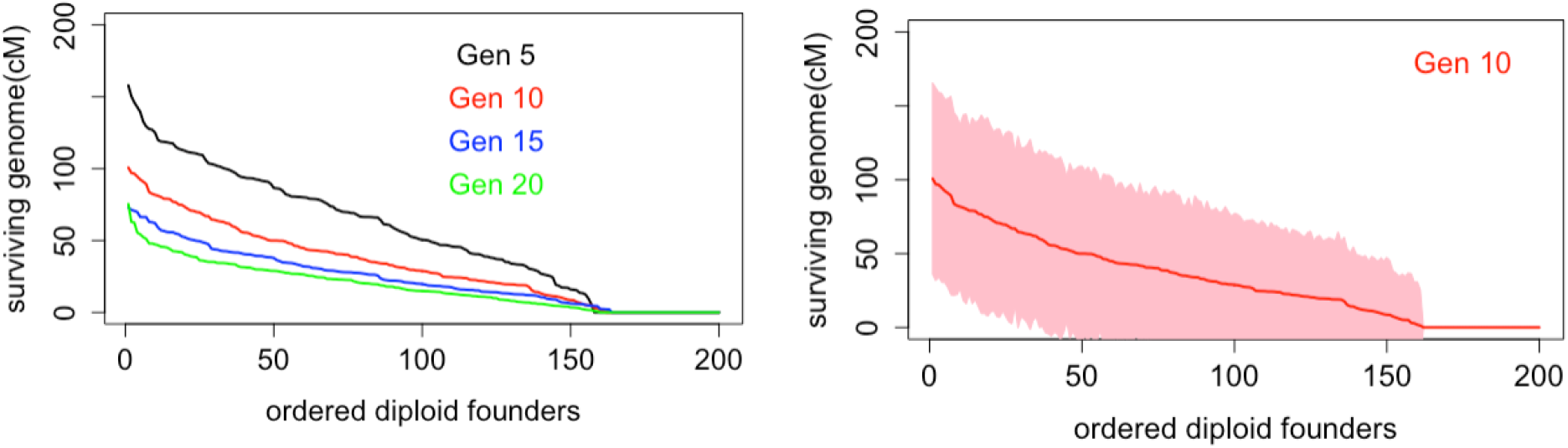
Survival amounts from a 1-Morgan diploid genome pairs; Left: mean values. Right: Variation at generation 10

However the expected proportion of genome surviving is simply the point-wise survival probability well-understood from single-locus analyses. Of much greater interest is the variation in genome survival. The right side of Figure 16 shows the variation at generation 10 over 500 replicates on a single population pedigree, shown as *±*2 empirical standard deviations conditional on non-zero surviving genome. The left side of Supplementary Figure C.1 shows the same for generations 5,10,15,20. We see considerable variation over repeated realizations on the same pedigree. The individuals with largest mean lengths of surviving genome of close to 100cM (50%) may have as little as 40cM or as much as 160cM surviving. At the other end of the scale, individuals with on average little surviving genome, may in some realizations have as much as 25cM surviving. Note that the variation over realizations in a given population pedigree is also the variation over different 1M chromosomes in a single realization of the population history.

Suppose however we are not interested in the amount of surviving genome for each individual, but only in the distribution across the 200 founders. That is, in any realization, the maximum amount, next and so on. The same realizations as before may be scored, ordering the values in each realization, with labelling of individual founders now lost. Figure 17 shows on the left the result for labelled founders as before, while on the right we have the anonymized population distribution at generation 10. Supplementary Figure C.1 shows the values for generations 5,10,15,20. Two things are apparent in Figure 17. First the variation over replicates is very significantly reduced. The mean maximum surviving genome at generation 10 is now about 150 cM, but this maximum will be attained by different founders in different realizations. Second, there are more founders whose pair of 1-Morgan genome lengths are extinct. In addition to the 40 individuals (20%) who have no descendants, in any realization an additional 20 diploids are likely to have both their 1-Morgan genome lengths extinct after 10 generations. The median among the 140 with some surviving length from the diploid 200cM is little changed (about 50 cM) but the standard deviation is about 5 times smaller.

**Figure 17:**
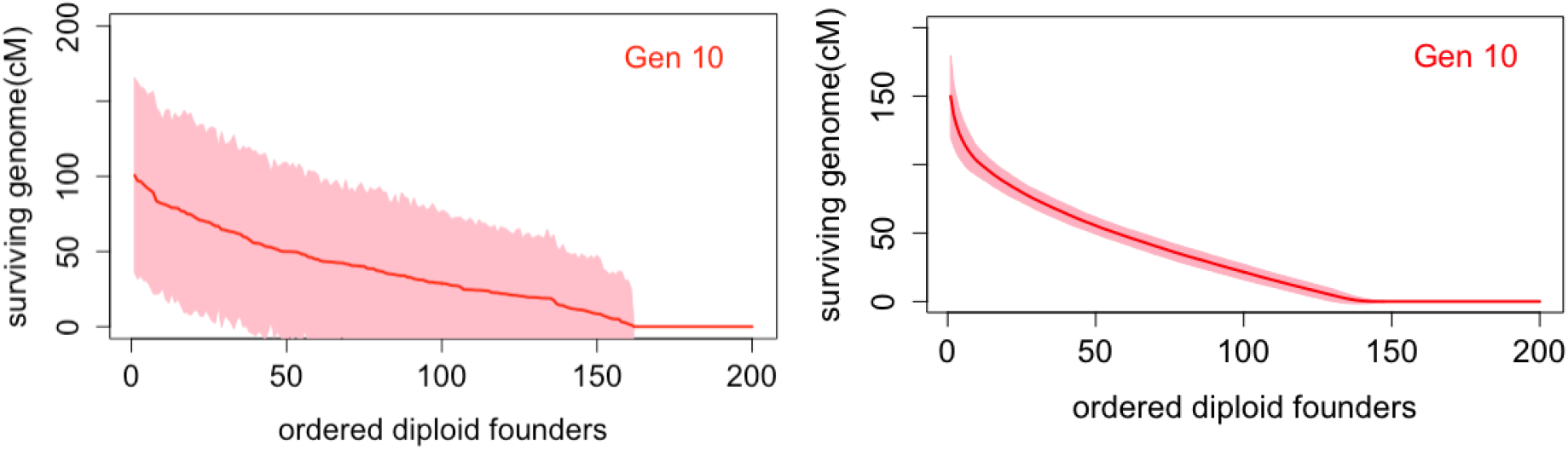
Survival amounts from a 1-Morgan diploid genome pairs; Left by founder individual. Right: By population distribution

#### 3.3.2 Variation within and between population pedigrees

Once individuals are anonymized, we can compared across population pedigrees. In Figure 18 on the left is shown again the results for 500 realizations on a single population pedigree, and on the right is one realization on each of 500 independently generated random-mating pedigrees. Supplementary Figure C.2 again gives the results for other generations, while Supplementary Figure C.3 gives additional discussion of results for the first few generations. The plot below in Figure 18 gives, plotted over the 20 generations, the mean over identified individuals of empirical standard deviations of:

**Figure 18:**
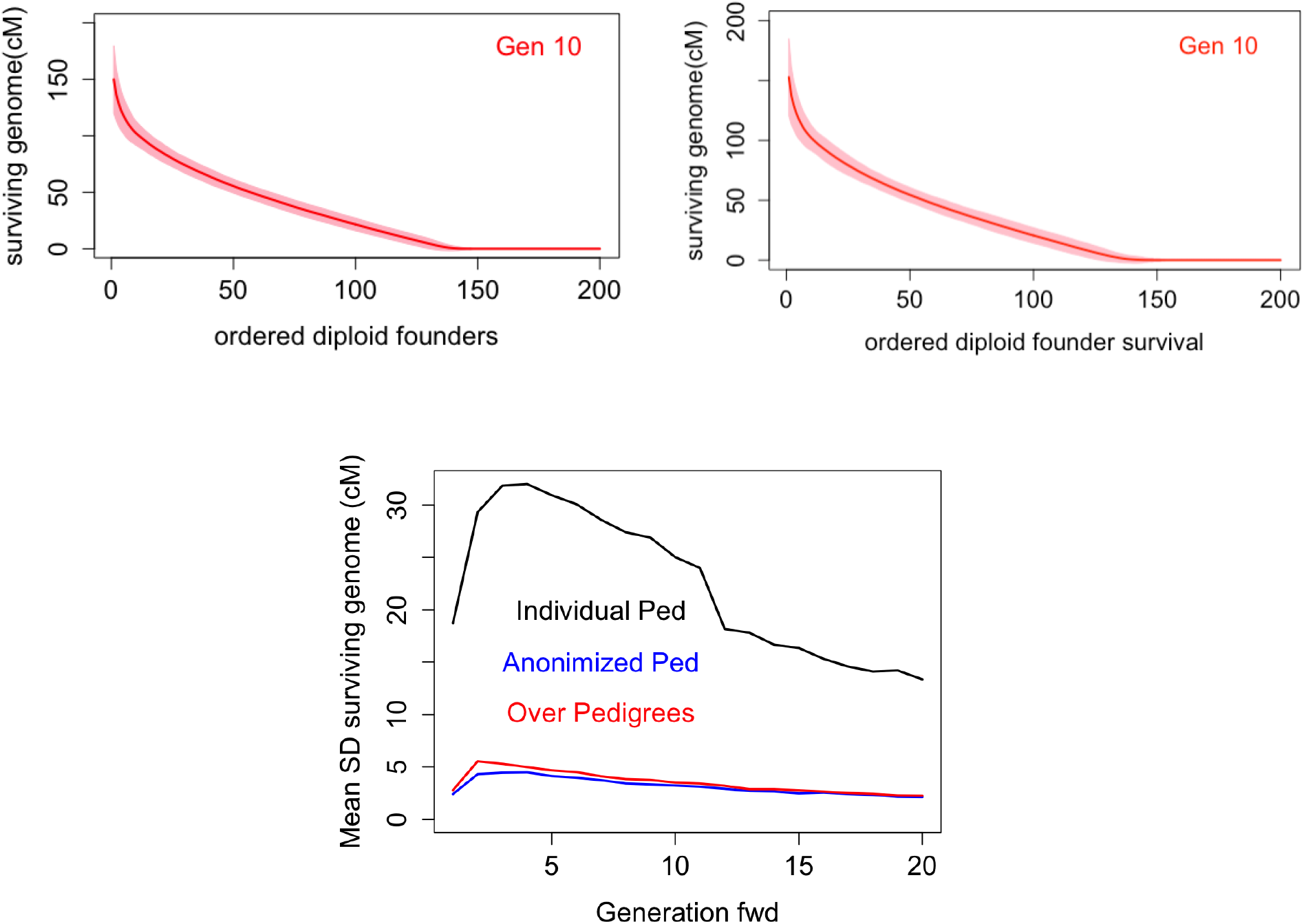
Comparison within and across pedigrees of survival amounts from a 1-Morgan diploid genome pairs. Above: single pedigree (left) and multiple pedigrees (right). Below: Mean empirical standard deviations of individual genome survival conditional on non-extinction (For details see text)

in black, 500 realizations on one pedigree;

in blue, the same 500 realizations on one pedigree with the individuals anonymized;

in red, one realization on each of 500 randomly generated pedigrees.

The variation across pedigrees is slightly larger, but the effect is scarcely apparent after the first few generation. The variation is in the meiosis pattern on the pedigrees, not among pedigrees generated with the same population model.

#### 3.3.3 Maximal descendant counts

We repeat the same sequence of operations for a rather different statistic, the maximum number of descendants of each haploid founder genome (FGL) across the 1 Morgan length, again using the approach of Section 2.4. The results are shown for generation 10 in Figure 19, and for other generations in Supplementary Figure C.4. In this case the FGL are ordered by mean total descent, which is a different ordering from mean maximal descendants, accounting for the slightly non-monotone mean ordering. In the left graph of Figure 19, similarly to the previous example, we see large variation in 500 realizations on a single population pedigree. Just about 20% of founder 1 Morgan lengths of genome are extinct in every realization, but conditional on some genome surviving, the variance over realizations is high. The group of FGL with highest expected total descent will have an average maximum of 15 to 20 descendants at some point in the 1-Morgan length, but will have 30 to 35 in some realizations. The median maximum among those FGL with descendants is about 10, with a large number of FGL showing means around 10. However, in some realizations these FGL may have as many as 25 descendants at some location.

**Figure 19:**
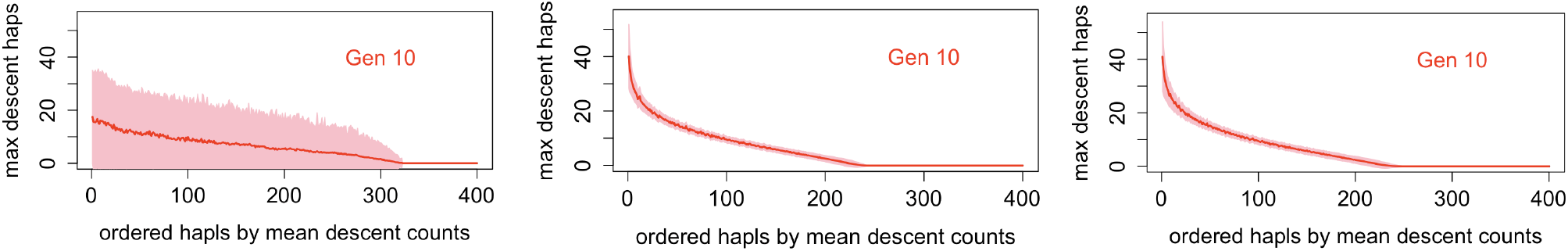
Maximal descendant counts of founder haploids (FGL) across 1-Morgan. Left: Single pedigree by founder haploid. Center: Single pedigree by populations distribution. Right: Across pedigrees

However, the anonymized version of the population distribution of the statistics has far less variance (Figure 19: Center). In most realizations there will be some founder genome that has 40 copies at some point in the 1-Morgan length 10 generations later, and in some realization there may be as many as 50 descendant copies. However, over most of the distribution, variation is very small, with a standard deviation of typically 2 or 3 copies. We see again the increase in extinction counts: whereas about 80 genomes (20%) are always extinct, in any realization on the pedigree a total of almost 160 (twice as many) is typically extinct. Recall the extinction of a 1-Morgan length in 10 generations an unbounded population is around 40% (Figure 6).

Again we may compare this distribution over founder genomes resulting from 500 realizations on a single pedigree, with 500 realizations each on a different randomly generated random-mating pedigree (Figure 19: Right). Variation is visibly increased, but overall the figures are very similar. From these and other examples we conclude that variation is in the meiosis process, rather than the pedigree process.

### 3.4 Dependence in survival of founder genome

In this section we return to the full 30M genome consisting of 30 1-Morgan chromosomes (Section 3.2), and consider dependence in survival and descent in a random-mating population of 100 males and 100 females each generation. Whereas at a single genome location, survival of one FGL can only negatively impact the survival of another, across the genome the reverse is true (Thompson, 2023). In an unbounded population there is strong positive dependence between the survival to a current genome of the two genomes (FGL) within any founder, due to junctions formed by recombination events in the first generation, at a rate 1 per Morgan in each meiosis. There is lesser but still strong dependence between the survivals of FGL in founder mates to a single current genome, for a similar reason.

The same pattern holds in our finite population, for survival in any current genome. While overall the dependence in descent among founders is slight (Figure 11) there is strong dependence between the two haploid genomes (FGL) of each diploid founder. Figure 20 shows scatterplots of the numbers of current genomes receiving descent from each founder’s maternal and paternal FGL at 10, 40 and 100 generations ago. The correlation is not only very high (0.9 to 0.85) but is maintained over the generations (Table 1). This results from multiple junctions formed between the two genomes at the very first generation–on average *L* = 30 per offspring. These junctions descend as single location markers, carrying the two genomes together. While many are lost, a few survive, and become frequent in the descendants. Descent may also be measured by the number of descendant segments of each founder genome, or, for a distribution with less less extreme tails, by the square root of this number. A figure showing dependence between a founder’s two FGL for this latter measure is given in Supplementary Figure D.1.

**Figure 20:**
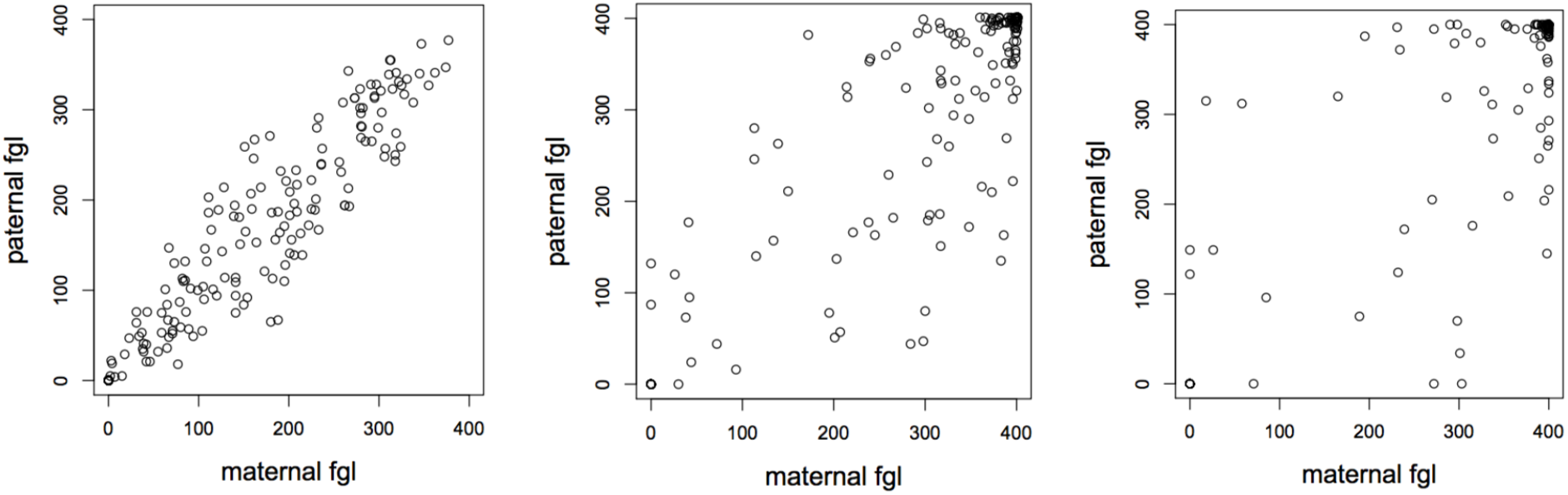
Number of genomes having descent from a founder’s maternal and paternal genomes in a random-mating population of 100 males and 100 females. Left: Generation 10. Center: Generation 40. Right: Generation 100.

**Table 1:** Correlations in descent of genomes from founders in a random mating population of 100 males and 100 females. Shown are correlations in descent between the two FGL of each founder, and correlations in descent from either FGL of founders who are founder mates.

|  | Generation 10 | Generation 40 | Generation 100 |
|---|---|---|---|
| Descendant genomes |  |  |  |
| within each founder | 0.9454 | 0.9140 | 0.9107 |
| in founder mates | 0.3541 | 0.3457 | 0.3354 |
| Descendant segments |  |  |  |
| within each founder | 0.9008 | 0.8606 | 0.8151 |
| in founder mates | 0.2383 | 0.2275 | 0.2418 |
| Sqrt descendant segments |  |  |  |
| within each founder | 0.9434 | 0.9067 | 0.8773 |
| in founder mates | 0.3105 | 0.2910 | 0.2991 |

Supplementary Figure D.2 gives a similar scatterplot to Figure 20 this time for the counts of genomes that are genetic descendants of either FGL of founder mates. Empirical numerical correlations between the two FGL within a founder and for FGL in founder mates are shown in Table 1. In a random-mating population high correlations do not persist beyond the first generation. While there is a positive correlation, it is far lower — around 0.34 for descendant genomes or even lower for the descendent segment measures. However it is again stable over time (Table 1). This correlation for founder mates results from junctions between their genomes formed in meioses from their offspring to the next generation. However, in a random-mating population an individual with many offspring will have many different mates; these mates may have few offspring, or many but with other mates.

## 4 Extensions to the population model

In this section we consider the effects of varying population size (Section 4.1), of mating structure (Section 4.2), and of family-size distribution (Section 4.3).

### 4.1 Population size

#### 4.1.1 Individual extinction in a very small population

In Section 3.1 we saw that population size has little effect on extinction of individuals, or on the total length of surviving genome from individuals founders.. However, at the extremes of the smallest populations or longest times, there are, of course differences. In the unbounded population, each single-location FGL becomes extinct with probability 1, although the expected time to extinction is infinite. In a finite population the ultimate extinction probability is (1 *−* 1*/*4*N*) since some FGL must survive at each location. In a population of 1 male and 1 female (*N* = 1, repeated full-sib-mating), both founders have descendants for all time. The case *N* = 2 is also easily dealt with: denote the individual extinction probability by *Q*. In any population, once a founder is an ancestor either of all males or of all females at some subsequent generation, they are immediately ancestors of all individuals at subsequent generations and can no longer become extinct. Thus if *N* = 2 there are only two transient cases. Either the founder has a single descendant (either male of female), in which case the probability of its future extinction is the same as the initial probability *Q*, or the founder’s descendants consist of 1 male and 1 female, in which case the probability is *Q*^2^, by the random mating assumption. Hence

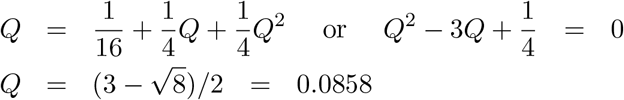

For larger values of *N* simulation is the simpler approach. For small *N* (e.g. *N* = 5 or *N* = 10) ancestral simulation using the parental probabilities of Supplementary Equation (A.1) is straightforward. For larger *N*, direct population simulation of ancestry or descent is simpler. In either case 10^6^ realizations can be rapidly performed. Figure 21 shows the individual extinction probability at 40 generations (effectively the ultimate value) for *N* = 1, 2, 5, 10, 20, 40, and 100. Simulation results confirm the value 0.086 for *N* = 2 and the unbounded-population value 0.203 for *N* = 100. although the value in a population less than half the size (*N* = 40, 2*N* = 80) is almost the same. The probability that a diploid individual has no long-term descendants is significantly smaller only for the very smallest populations.

**Figure 21:**
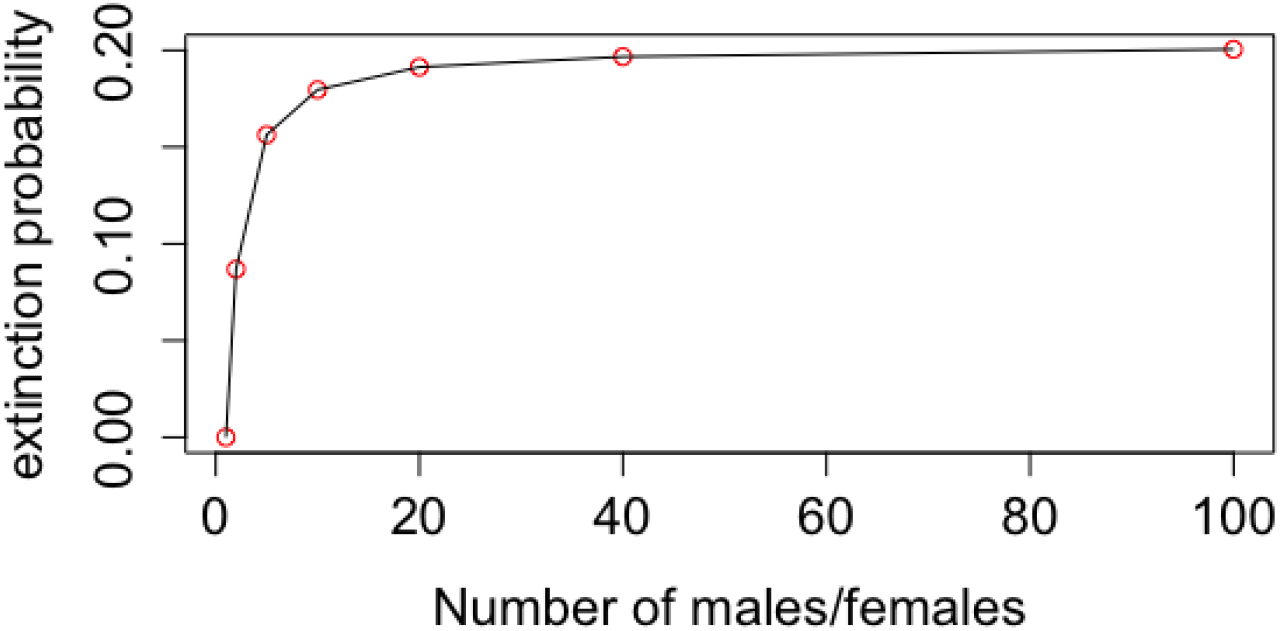
Probabilities of individual extinction in 40 generations in diploid random-mating populations with *N* males and *N* females in each generation, for *N* = 1, 2, 5,10, 20, 40, and 100.

#### 4.1.2 Descent and ancestry in a population size 2000 over 100 generations

For comparison with the results of Section 3.2 we simulated the descent of genomes of *L* = 30 Morgans, in a random mating population of 1000 males and 1000 females per generation over 100 generations. The population was simulated downwards, using the *ibd create* program of MORGAN Ver 3.4 (2017). The descent of the 4000 founder genomes was simulated, and founder genomes (FGL) and breakpoints were output output for each of the 4000 genomes of the final generation. Here we summarize results of FGL segments disregarding the segment lengths determined by the breakpoints.

Overall 877 (21.9%) of founder genomes (FGL) are extinct, and 3123 survive in at least one current individual. Consider first the current genomes. The number of founder segments in each current genome is approximately Normal, and has mean 2973 and standard deviation 54.7; these are very close to the expected values for the Poisson *Po*(30 *×* 100). The number of FGL present in each current genome is also approximately Normal with a mean of just under 1600, and a standard deviation of 23. Unsurprisingly the count of segments and count of FGL are correlated (Correlation 0.65), but there is no tight relationship.

By contrast, Figure 22 shows the results from the perspective of founder genomes (FGL). On the left is shown the total number of copies of each FGL present as segments in current genomes, ordered by their number of such copies. Note 877 are extinct (0 current copies). As in the smaller population (Section 3.2.1). the median count of descendant segments is almost exactly 2000 (See Figure 12 (Left)). That is 2000 FGL have 2000 or more current descendant segments. There are 3 FGL that are represented by more than 20,000 segments; of course many of these are duplicate and/or overlapping segments present in multiple current individuals. In the center is plotted the number of descendants in which each FGL is present. On the right is a scatterplot of the number of descendants against the number of segments: the two are quite closely related. Of course with 0 descendant segments, there are 0 descendants, but the tight relationship is surprising. FGL with more than about 12,000 descendant segments are present in almost every current genome, but only 93 (2.3%) of the 4000 FGL have this many descendant segments. To compare with the 10 times smaller population (Section 3.2.1), there FGL with at least 1200 segments were present in every current genome, but by 100 generations about 50% of FGL achieve this.

**Figure 22:**
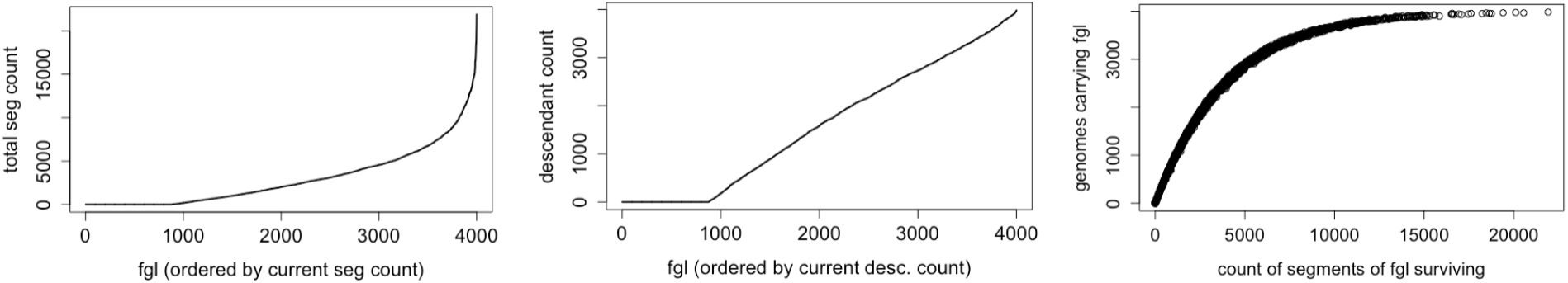
Counts of FGL in acurrent 2000-member population after 100 generations. Left: Count of founder segments. Center: Count of distinct FGL. Right: Scatterplot of the number of FGL vs number of segments.

Population size does affect any population statistic that is a function of random genetic drift. In particular it affects the rate of change of FGL frequencies for those FGL that are present in multiple copies. The number of current descendant genomes in which each FGL is present is such a statistic. In the center plot of Figure 22 we see a surprisingly linear patters for the ordered numbers of descendant genomes is which each FGL is present: recall we saw the same result at 10 generations in the 10 times smaller population (Section 3.2.1). Overall, with appropriate scaling of time, descent is little affected by populations size.

#### 4.1.3 Dependence in survival of founder genomes

In this same single population of 1000 males and 1000 females, corresponding to the results of Sectiion 3.4 for populations of 100 males and 100 females, we consider correlations at 100 generations in the descent from the two FGL of each founder, and in descent from either FGL of individuals who are founder mates. Supplementary Figure D.3 shows the scatterplots and the correlations are given in Table 2. We see the correlations in counts of descendant segments are comparable to those of the 10-times smaller population (Table 1), but those for the number of genetic descendants are slightly lower. In a very large (unbounded) population, the only way in which the 2 FGL of a funder will coexist in a descendant is by survival of a junction formed at the first generation. In a smaller population the ultimate survival of such a junction (1*/*4*N*) is slightly higher, but also there is higher probability that two specific FGL will coexist in a descendant due to mating of relatives. Note also that in large populations there are a few FGL with extremely large numbers of descendant segments; these distort correlation measures.

**Table 2:** Correlations in counts of genomes that are genetic descendants and of descendant segments.

|  | FGL within founders | FGL in founder mates |
|---|---|---|
| Number of genomes with genetic descent | 0.860 | 0.259 |
| Total number of descendant genetic segments | 0.794 | 0.214 |
| $\sqrt{\text{Total number of descendant genetic segments}}$ | 0.875 | 0.254 |

### 4.2 The effects of mating structure

So far we have considered only random mating populations. In Table 3 below are shown three alternative forms of random mating with varying structures of mating couples. The first line of the table corresponds to true random mating in a dioecious population Each of 2*N* offspring is assigned a random father and a random mother, each sampled with replacement from the parental males and females. In the second case, each of *N* male offspring is assigned a random father and a random mother, each sampled with replacement for the parental males and females, and a female offspring is then assigned to the same parent couple. This creates pairs of full siblings with larger sets of half-siblings. In bottom row of the table., *N* couples are randomly formed, but then fixed. Each of 2*N* offspring is independently assigned to a couple. In this case, all offspring fall into full sibships. We again consider a genome of length *L* = 30 Morgans.

**Table 3:** Correlations in descendant counts of genomes from founder mates in populations of 100 males and 100 females.

| Offspring assignment to<br>to parent couples | Number of<br>descendant genomes | Number of<br>descendant segments |
|---|---|---|
| 2*N* random couples; with replacement<br>Each offspring randomly assigned | 0.25 to 0.28 | 0.16 to 0.29 |
| *N* random couples ; with replacement<br>1 son, 1 daughter each | 0.33 to 0.35 | 0.22 to 0.24 |
| *N* couples sampled; no replacement<br>2*N* offspring independently assigned to a couple | 0.85 to 0.95 | 0.83 to 0.92 |

Note that in the first and third cases the marginal offspring distribution for each parent is the same, since each of the 2*N* offspring is independently assigned to parents.. The offspring number is *Bin*(2*N,* 1*/N*) with mean 2 and variance 2(1 *−* 1*/N*), or effectively Poisson mean 2 unless *N* is very small. The probability of no offspring is 0.135 and of a single offspring 0.271. The probability of more than 6 offspring is 0.005. In contrast, the offspring distribution in the second example is 2 *× B*(*N,* 1*/N*). so again having mean 2, but variance 4(1 *−* 1*/N*). Unless *N* is small, it is effectively twice a Poisson mean 1, with probability 0.36 of no offspring, and a probability 0.019 of more than 6 offspring. Other examples of the second mating structure in which the variance in offspring count is varied, including an extreme example in which the variance in offspring count is reduced to zero, are considered in Section 4.3 and Supplementary Section E below.

Marginal statistics for extinctions and for counts of genetic descendants and descent of genome segments are indistinguishable for the first and third scenarios with *N* = 100, and surprisingly little changed in the second despite its greater variance in offspring number (Results not shown). Nor does mating structure impact correlations between the two genomes of each founder: these remain at 0.85 to 0.95 for the count of distinct descendant genomes, and 0.8 to 0.9 for the count of descendant genome segments. However, the form of mating structure does have a big impact on dependence between founder mates in their counts of distinct descendant genomes, and of descendant genome segments (Thompson, 2025).

Table 3 shows the correlations in descent from founder mates under the three different forms of random mating in the population of 100 males and 100 females at each generation. Shown are both correlations in the number of descendant genomes in which either genome of each founder genome is represented, and the total number of genome segments descending from either genome of each founder. The numbers shown are approximate from a small number of realizations. They are broadly constant over time, since they derive from junctions formed in meioses of the immediate offspring of founders. The random mating results (first line) are those already seen in the Section 3.4 (Table 1 and Supplementary Figure D.2). Despite the creation of full siblings and larger sets of half-siblings, the second row shows only about a 25% increase in the correlations between founder mates. In contrast is the bottom row of the table, in which monogamy is enforced. The full sibships restore the correlations between mates to values comparable to those seen between the two genomes of each founder. In this scenario, if a diploid founder has many offspring so does his/her mate, and at the next generation many junctions will be formed between the pairs of haploid genomes of the founder mates.

### 4.3 Variance in reproductive success

We briefly consider two examples of the effect on individual and genome extinction of two versions of variation in reproductive success. The first is a version of a random mating population in which 2*N* offspring are assigned at random to females and male parents. The second involves the formation of couples, with each mating producing a male and a female offspring. In all cases the population will have *N* females and *N* males at each generation, and the examples will use *N* = 100. Both sampling schemes ensure that, marginally, all individuals are equivalent; each individual has an expected 2 offspring.

#### 4.3.1 Variance in male reproductive success

The population pedigree is formed by randomly assigning each offspring a male and a female parent. The female parent is assigned independently for each offspring with probability 1*/N* for each potential mother, as in a random mating population. Male parents are assigned sequentially. When *M* male parents have been assigned, a male who has already been assigned *m* offspring will be assigned the next with probability (*βm* + 1)*/*(*βM* + *N*). The probability 2 offspring have the same mother is 1*/N*, the effective number of mothers is *N*, and the variance of offspring number is the binomial *Bin*(2*N,* 1*/N*) variance 2(1 *−* 1*/N*). For the fathers, the probability 2 offspring have the same father is (1 + *β*)*/*(*N* + *β*), by the exchangeability of orderings of fathers under the Polya-type weighting scheme. Thus the effective number of fathers is (*N* + *β*)*/*(1 + *β*), and the overall effective populations size is

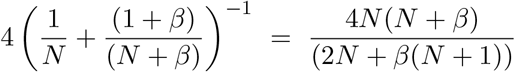

Although we could consider varying *β* and *N* to compare populations with the same effective size, recall that changes in *N* has almost no impact on individual extinction and on the proportion of genome surviving (Sections 3.1 and 4.1.1). We have therefore preferred to consider the effect on variance in offspring number as *β* changes.

The variance of male family size can be derived as follows. If family size *X* has *P* (*X* = *m*) = *p_m_* the probability that an offspring comes from a family size *m* is *mp_m_/E*(*X*). With *E*(*X*) = 2 the probability two offspring have the same father is then

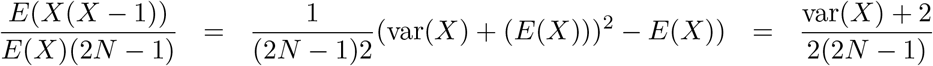

Setting this equal to (1 + *β*)*/*(*N* + *β*) we obtain

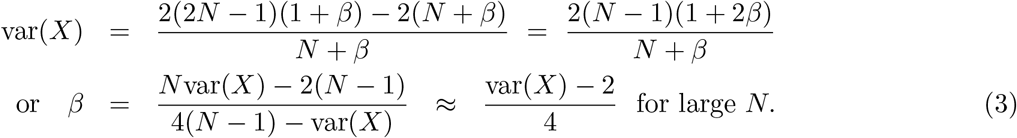

These formulae are in accord with simulation results. For *N* = 100, *β* is essentially linear in var(*X*).

For *β >* 0, the standard deviation of *X* is greater than the Binomial 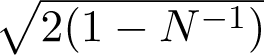. For a standard deviation of male offspring number 2.5 (variance 6.25) we have *β* = 1.0956. Then both males and females have mean offspring number 2, and the females have variance just under 2, so the overall variance is about 4. Using this value of *β*, Figure 23 shows on the left the probability of extinction for males in blue and females in red, with the values for a random-mating population shown as a dashed line for comparison. Unsurprisingly, the extinction probability is greatly increased for male diploid founders. It is also substantially increased for female founders: fathers have daughters and mothers have sons. On the right is shown the expected proportion of genome that is extinct. Here the effect is much less marked, although both males and females receive an initial hit, the males especially so due to the large first-generation probability of zero offspring. The difference between males and females rapidly declines. The difference relative to the random mating population persists, since survival is largely determined by the initial generations. The difference eventually declines to 0 since the ultimate expected proportion is (1 *−* 1*/*4*N*) when only 1 genome survives at any point in the genome, but the difference is still clearly apparent at 100 generations.

**Figure 23:**
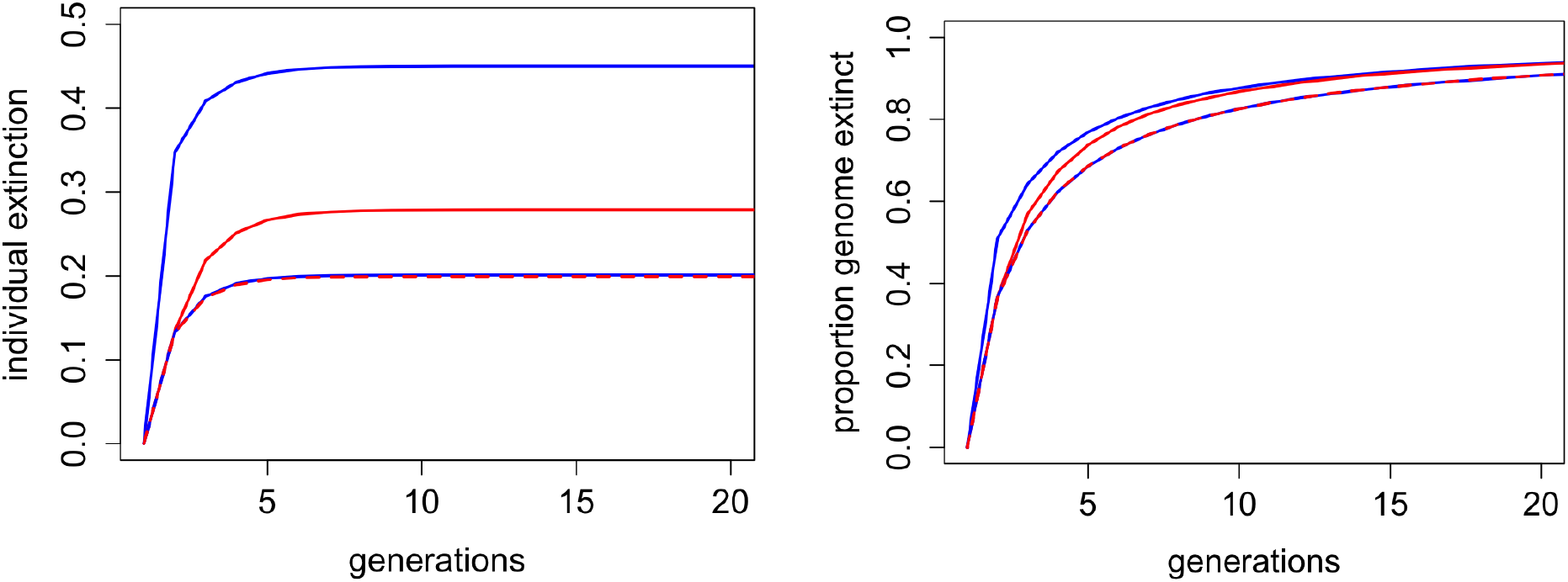
Comparison of *β* = 1.0956 with random mating, with males in blue and females in red. Left: diploid extinction probabilities. Right: Expected proportion of genome extinct. (The dashed lines show the random-mating result: the same for males and females).

Figure 24 shows on the left the mean number of descendants conditional on survival, while the right shows the descendant counts for the 100 males (blue) and 100 females (red) for one realization. Apart from a small boost for males in the early generation there is little effect of the mean descendant counts conditional on survival, and the time for all non-extinct individuals to achieve ancestry of the entire population (16 or 17 generations) scarcely affected. The righthand figure may be compared with the righthand figure of Figure 10 for a random-mating population. The variance among individuals is slightly higher, with a few males achieving ancestry of the entire population quite rapidly, but again the effect is not marked.

**Figure 24:**
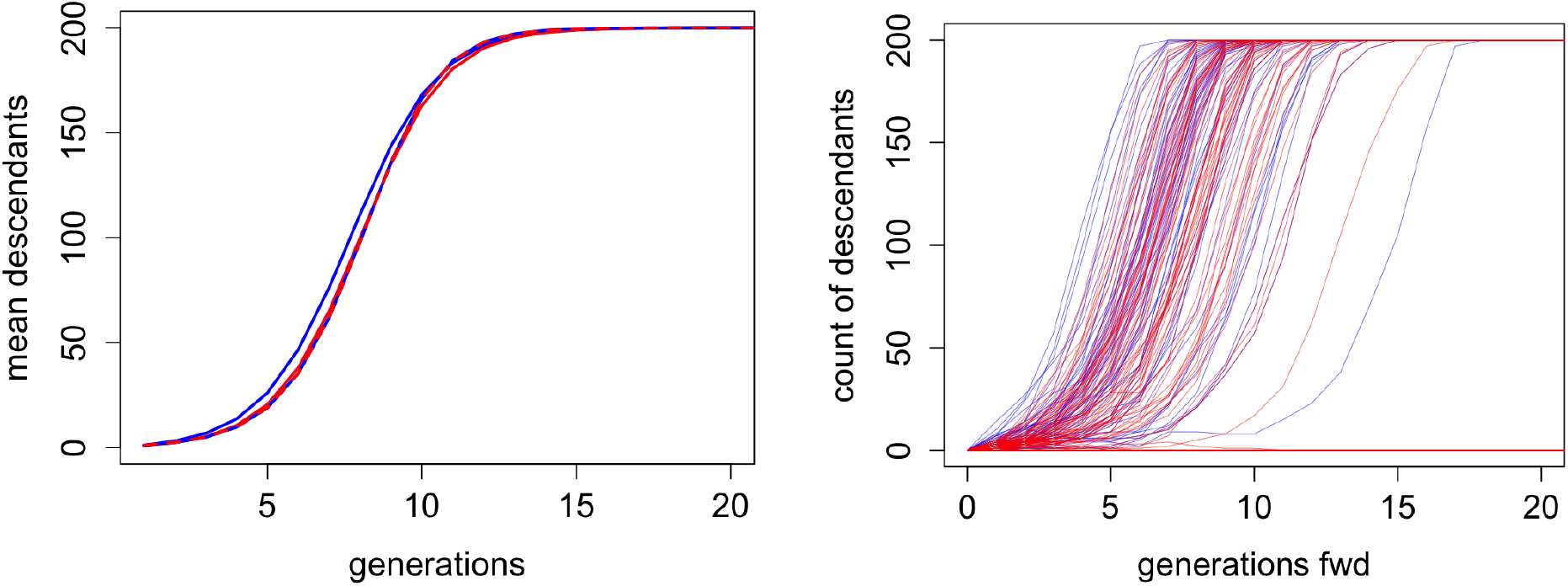
Comparison of *β* = 1.0956 with random mating, with males in blue and females in red. Left: mean number of descendants conditional on non-extinction: the dashed line shows the random mating result. Right: One realization of descent.

#### 4.3.2 Variance in re-mating probabilities

A visiting student Zheng Cai (Pers.comm., 2009) wishing to produce population pedigrees with equal numbers of males and females, constructed matings in which each produced one son and one daughter. If males and female parents are sampled with replacement we have the scheme already discussed as the middle example of Table 3. The variance of offspring number is approximately 4. If male and female parents are sampled without replacement, each parent is sampled exactly once, and has one som and one daughter. The probability of individual extinction is 0, as is the variance in offspring number. While this scheme is highly unrealistic it has two advantages. First it produces pedigrees is which ancestry and descent are equivalent. Every parent has one son and one daughter. Every offspring has one father and one mother. Second, as the maximally constrained pedigree, it serves to demonstrate the limits on the constraints of pedigree on genome survival and descent. Supplementary Figure E.1 shows the equivalence of ancestry and descent, and the reduction in variance both among individuals within a pedigree, and over pedigrees. As previously, there is scarcely any difference in the pattern among individuals within a pedigree compared to individuals among pedigrees, and the overall pattern is very similar to other results. The main effect is. the slightly reduced time to complete ancestry and descent, with completion by 12 generations rather than the value of about 16 or 17 for a random-mating population of 100 males and 100 females.

Grimson (2016) extended previous work on population pedigrees and modified Cai’s scheme to produce values of offspring number variance between 0 and 4. Matings are formed sequentially by choice of a random male and a random female, and a son and a daughter is assigned to each mating. When parents in *M* matings have been assigned, with *m_j_*assigned to male *j* (Σ*_jj_ m_j_* = *M*), the male parent *i* for the next mating is selected with probability *α^m^^i^/*(*N*Σ*_jj_ α^m^^j^*), and so also for female parents. That is, each assigned mating down-weights the parents by a factor *α* for subsequent matings. Note *α* = 0 corresponds to sampling without replacement and variance 0, while *α* = 1 corresponds to sampling with replacement, with the variance of mating number approximately equal to 1, and of offspring number approximately equal to 4, for large *N* . Unfortunately, orderings of selections are not exchangeable under this scheme, while the previous additive Polya-type scheme with negative *β* risks hitting the 0 bound for mating probability and also thence becoming non-exchangeable. Thus explicit formula such as Equation (3) are not available, By simulation, Grimson (2016) determined a value *α* = 0.33 for her large-population study. For a population of *N* = 100 males and 100 females, we find *α* = 0.383 gives a variance of mating number almost exactly 0.5, and hence a variance of offspring number 2. (In the range 0.3 to 0.4, the variance is relatively insensitive to *α*.)

Figures 25 and 26 show the results for *α* = 0, 0.383, and 1, for the same statistics as in Figures 23 and 24. Note *α* = 0.383 is chosen to given the random-mating variance of offspring number, while *α* = 1 gives a variance of 4, comparable to *β* = 1.0958 of the previous figures. For *α* = 1 the extinction probability is comparable to the value for males under the previous scheme, while *α* = 0.383 gives higher probability than does random mating. The expected proportions of genome extinct are surprisingly close the the previous values for these two curves. The case *α* = 0 with diploid extinction probability 0 has a lower expected proportion of of extinct genome. Figure 26 shows an interesting feature. While conditional on non-extinction the higher *α* values show high mean number of descendants, the zero-extinction curve *α* = 0 overtakes these at about 10 generations, and reaches complete ancestry (200 descendants) by about 12 generations. (See also Supplementary Figure E.1.) The righthand figure for *α* = 0.383 may again be compared with the random-mating case Figure 10; the two figures are very comparable, with complete ancestry being achieved by survivors between 7 and 14 generations. Note these two right-hand figures each relate to a single realization of a population pedigree: some variation is expected.

**Figure 25:**
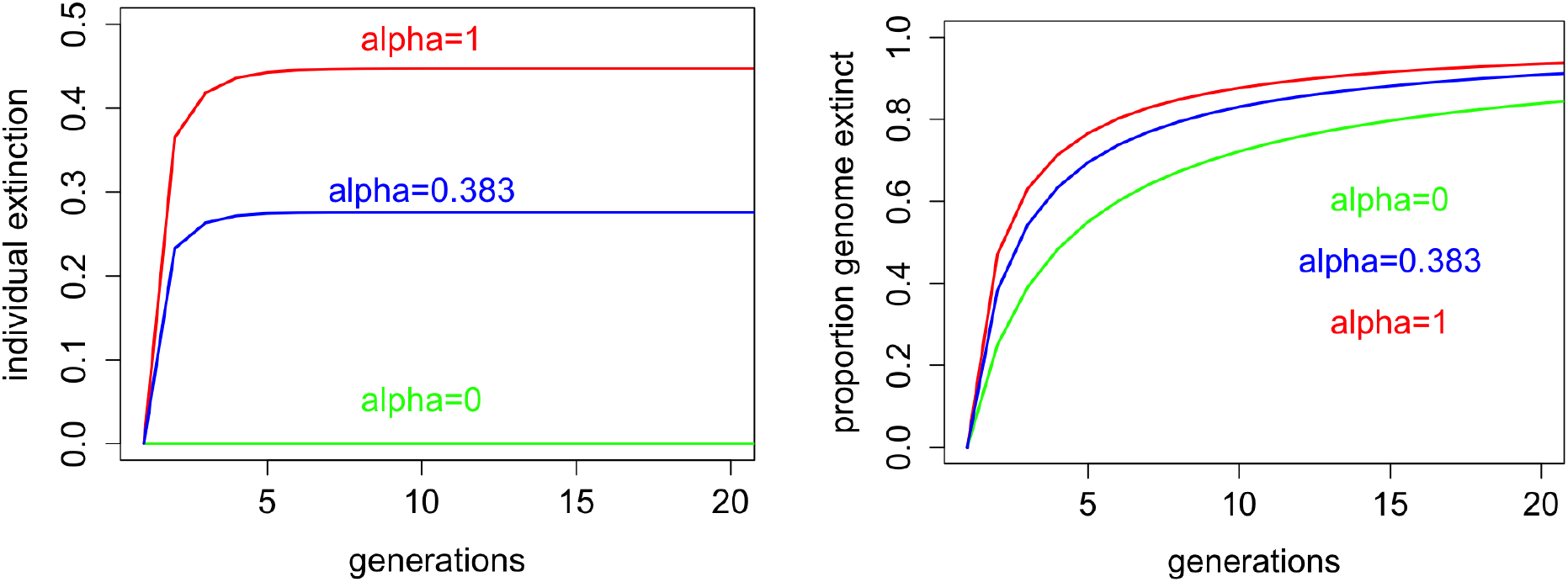
Comparison of varying re-mating probabilities for variance of offspring number 0,2,4: Left: diploid extinction probabilities. Right: Expected proportion of genome extinct.

**Figure 26:**
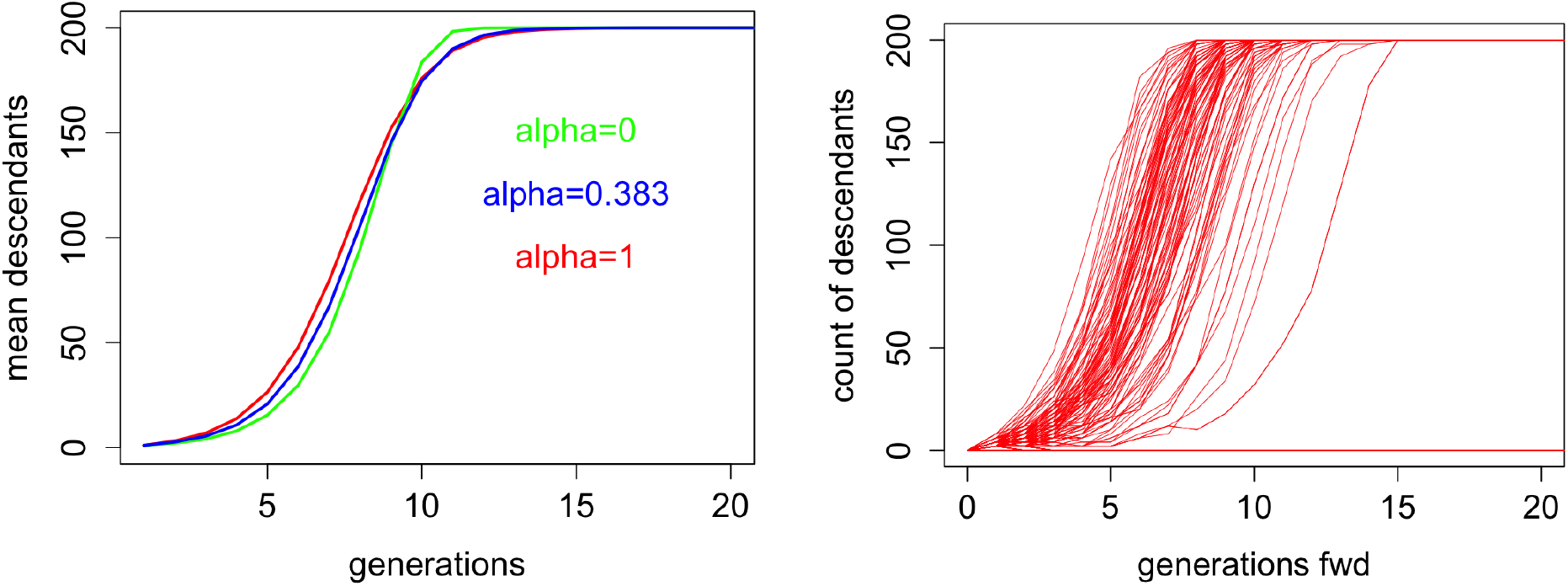
Comparison of varying re-mating probabilities, for variance of offspring number 0,2,4: Left: mean number of descendants conditional on non-extinction.. Right: One realization of descent

## 5 Discussion

This paper has investigated many aspects of genome survival and descent in a small diploid dioecious population, taking as its main canonical example a random-mating population with 100 males and 100 females in each generation. Typically we have considered a (haploid) genome length of 30 Morgans, although results for Section 2.4 we focus on a 1-Morgan length of genome typical of a mid-size mammalian chromosome. We also consider descent from the diploid founders, which provides a limit for an infinite genome, and a framework within which finite founder genomes segment and descend and disperse through the population. We consider only the Haldane meiosis model of no genetic interference. Under this model the division of the 30M genome into *C* chromosomes has only small effect, creating (*C −* 1) first-generation obligatory opportunities for ”recombination” each with 0.5 probability. As founder chromosomes become broken into segments, and segments dispersed among the members of the population, the effect on results and conclusions is minimal. An exception may be the very smallest chromosomes with genetic length 0.5 Morgans, due to the requirement of at least 1 chiasma on the tetrad for the meiosis process, but having relatively much smaller physical length. Considering the effects of the Haldane model versus a complete interference model remains a topic for the future.

Most results are by simulation, and are designed to be sufficient to be illustrative rather than definitive, typically of 500 or 1000 realization of events on the population pedigree. Simulations were carried out to a depth of 100 generations, but most results are presented only to 20 or 40 generations: in a small population the stage for survival and descent is typically set within 20 generations, in terms of counts of descendants, and rank within the population for surviving genome and segment descent copies. Although this paper focuses on descent and survival of genome, all simulations may in principle be carried out either forwards or backwards in time. The population pedigree determines the existence of parent and offspring links, and. meiosis breakpoints occur independently in every meiosis (except that monozygous twins share their parental meiosis). At any point in the genome, any ancestral lineage can be traced upwards from current (haploid) genome to a unique haploid genome (FGL) of some diploid founder, Likewise the lineage may be traced downwards from FGL to current genomes, although of course the FGL may have multiple descendant copies at any point in the genome. Both upwards and downwards simulations are used in this paper, but for many purposes upward simulation is more efficient, since only segments of ancestral genome surviving in the current population are relevant. A key methodological development is in Section 2.4 where it is shown that previous methods of simulation of ancestry of a single haploid genome (Thompson, 2024), can be used to simulate a variety of statistics of founder genome survival and descent. This approach is used in Section 3.3.

One result that emerges in several contexts is the short time-scale of relevant events. The number of offspring and grand-offspring in the first two generations has a major impact on survival of genome and on descendant copies of that genome. Within about 17 generations a diploid founder is an ancestor either of the entire population or of nobody. Genome spreads a little more slowly, but the large majority of diploid individuals with descendants have their genome represented in almost all individuals within 25 generations (Figure 13, and related results). However, there are a few individuals with few early-generation descendants who nonetheless become ancestral to the entire population. For these individuals much of their genome is lost; their descendant segments largely overlap and are present in a more limited proportion of individuals.

In terms of dependence in survival and descent among founder genomes, the first generations are critical. There is strong positive dependence in the descent between the two genomes of each founder, and a lower but still positive dependence between the genomes of founder mates. Only enforced monogamy extends the high dependence a single generation to founder mates. Once segments of each FGL are dispersed in the population, any dependence decays rapidly. There is also, unsurprisingly, positive dependence of the length of a founder genome surviving, and the total descendant length of that founder genome (Figure 15). There is also high dependence between the number of segments of descent from a founder, and the number of descendants that founder’s genome is represented in, although the latter becomes saturated within about 40 generations. (Figure 12). Of course, there is very high correlation between total descendant genome FGL length and total number of segments of descent of each founder genome FGL at any given generation.

Another key aspect is comparison of the variability among individuals within a population pedigree over realizations on that single population pedigree, and over different pedigrees generated according to the population process. These comparisons are facilitated by the methods of Section 2.4. Earlier results (Figures 10 and 11) had suggested that variability among the members of a population differed little from that of individuals independently generated by the population process. The results of Section 3.3 explore this in more detail for the rather different statistics of surviving diploid founder genome length and for the maximum number of descendants of any FGL across the length of genome considered. For these and other measures (not shown), the individual variation over realizations on a population pedigree is high, with standard deviations similar to the mean.. There is huge variation in meiosis, and in particular on outcomes resulting from the first very few generations of the pedigree.

However, while each FGL may vary in its success, the overall distribution of the statistic across the population is far less variable. That is, for example, the amount of genome surviving from each of the most successful diploid founders or FGL varies widely, but over realizations the ranked amounts surviving from each founder varies little. More surprisingly, the variation in the distribution over members of independently generated population pedigrees is only slightly greater than for multiple realizations on a single pedigree. This is less so for measures of the extremes: each population pedigree will have its particular extreme events, whether a founder with exceptionally many first and second-generation descendants, or a founder with very few early-generation descendants nonetheless surviving to be the ancestor of everybody but with almost no surviving genome and the genetic ancestor of few (Supplement Section B). However, overall variation is in the meioses, not in the pedigree.

Finally we have considered some variations to the basic theme of a random mating population of 100 males and 100 females. The very smallest populations show increased survival, but this is mainly because some ancestral lineage must survive at each point in the genome. Larger population size has little effect on individual genome survival and descent, nor on any event that is determined within a few generations from founding; it does have an effect on classical population genetic measures such as the longer term rates of change of point frequencies of FGL. Mating structure per se also has little effect, although monogamy greatly increases the correlations in descent and survival between founder mates. Family size variance has a major effect on individual (or entire genome) extinction, and varying male reproductive excess also affects the probabilities for females: fathers have daughters, and mothers have sons. However, conditional on survival the proportion of an individual’s genome surviving is not greatly affected, and the effect is transient. In terms of mean and within population variance of time to complete ancestry effects are also small with 16 to 17 generations being the norm. An exception are the ”Cai” pedigrees, in which every mating is monogamous and produces a son and a daughter. In this extreme case variation is reduced, and every founder is either extinct or the ancestor of everybody by about 12 generations.

There are several implications for endangered populations being maintained or reestablished from a small group of founder individuals. A major hurdle is that the major part of variation is in meiosis not in pedigree: the are limits to what pedigree-based management can achieve. However, a few conclusions can be noted. First, the first few generations are critical: genome once lost cannot be regained (Geyer et al., 1989). Second, variance of offspring number should be reduced to the extent possible, and monogamy has advantages provided founder mating pairs are carefully chosen (Geyer et al., 1993). Reduction of offspring variance to 0 is mostly infeasible, and enforcing offspring of opposite gender even more so. However in captive managed populations in which offspring are easily produced, but costly to rear and maintain as adults, selective culling could potentially be employed, at least once the population reaches capacity for management.

## Statements and declarations

No funding was received to assist with the preparation of this manuscript. The author has no relevant financial or non-financial interests to disclose. A.I. was not used knowingly or intentionally in any aspect of this paper.

## Acknowledgements

I am grateful to Dr. Joe Felsenstein and to Dr. Mary Kuhner for many helpful discussions on many aspects of this paper. I am grateful to Dr.Phil Green for the insight of Section 2.3 that in considering the non-overlapping segments of ancestry of a single current haploid genome present in genomes of ancestral individuals who share a parent, these segments can be independently inserted into the parental genome, due to the independence of meioses.

