## Supplementary Figures for "Genome survival and descent in population pedigrees"

### Supplementary Material

#### A Iterative formulae for counts of diploid ancestors

This section provides an alternative to the complex inclusion-exclusion approach of Feller (1950) (Equation (2)) for the number of mothers (or of fathers) $X$ of a given number of individuals $A$ in a random-mating population of $N$ females and $N$ males. With $N$ fixed, it proceeds iteratively over $X$ and $A$. Suppose that at some past time the number, $A$ of diploid ancestors of a current individual is $A = a$ $(1 \leq a \leq 2N)$. Suppose that the number of mothers of these $a$ individuals is $X$ $(1 \leq X \leq \min(N, a))$, and note that the probability distribution of the number of fathers is the same as that of mothers, and that the counts of mothers and fathers are independent. To obtain the probability $P(X = x \mid A = a)$ consider successively assigning the individuals to their mothers; let $p(a, x)$ be the probability that $a$ individuals have $x$ $(x \leq \min(N, a))$. When $x$ distinct mothers have been assigned, the probability an additional individual will be assigned to a current mother is $x/N$ and to a new distinct mother is $(1 - x/N)$ so that

$$p(a, x) \quad = \quad p((a-1), x)\frac{x}{N} \; + \; p((a-1), (x-1))\left(1 - \frac{(x-1)}{N}\right) \tag{A.1}$$

with $p(1, 1) = 1$.

The above iterative formula also provides that

$$\mathrm{E}((N-X)|A=a) \quad = \quad N \sum_{x=1}^{\min(N,a)} (1 - \frac{x}{N})p(a, x) \quad = \quad (1 - 1/N) \;\; \mathrm{E}((N-X)|A=(a-1))$$

or $\mathrm{E}(X|a) = N(1 - (1 - 1/N)^a)$, although this is more easily seen by noting the fact that each of the $N$ potential mothers is assigned no offspring is $(1 - 1/N)^a$. The conditional variance can be similarly obtained, using the fact that a specific two potential mothers are assigned no offspring is $(1 - 2/N)^a$ so that

$$\mathrm{var}(X \mid A = a) \quad = \quad N(1 - 1/N)^a(1 - (1 - 1/N)^a) \; + \; N(N-1)\left((1 - 2/N)^a - (1 - 1/N)^{2a}\right)$$

Table A.1 shows the distribution for the case $N = 4$, for values of $a$ from 1 to 8. Numerical checks show agreement with the Feller formula (Equation (2)).

Combining the independent mothers and fathers, each with the distribution of $X$, we can derive expressions for the mean and variance of $A_{k+1}$ the number of ancestors at $(k + 1)$ generations ago given the number $A_k$) at $k$ generations ago:

| a | $x=1$ | $x=2$ | $x=3$ | $x=4$ | $\mathrm{E}(X \mid A=a)$ | $\mathrm{var}(X \mid A=a)$ |
|---|---|---|---|---|---|---|
| 1 | 1 | 0 | 0 | 0 | 1 | 0 |
| 2 | 1/4 | 3/4 | 0 | 0 | (16-9)/4 =1.75 | 0.1875 |
| 3 | $1/4^2$ | $9/4^2$ | $6/4^2$ | 0 | (64-27)/16= 2.32 | 0.3398 |
| 4 | $1/4^3$ | $21/4^3$ | $36/4^3$ | $6/4^3$ | (256-81)/64 = 2.73 | 0.4138 |
| 5 | $1/4^4$ | $45/4^4$ | $150/4^4$ | $60/4^4$ | $781/4^4 = 3.05$ | 0.4230 |
| 6 | $1/4^5$ | $93/4^5$ | $540/4^5$ | $390/4^5$ | $3367/4^5 = 3.28$ | 0.3926 |
| 7 | $1/4^6$ | $189/4^6$ | $1806/4^6$ | $2100/4^6$ | 14197/486 = 3.47 | 0.3426 |
| 8 | $1/4^7$ | $381/4^7$ | $5796/4^7$ | $10296/4^7$ | $58957/4^7 = 3.60$ | 0.2870 |

Table A.1: Probabilities that $a$ offspring have $x$ distinct mothers, for $N = 4$, and $a = 1, ..., 8$.

$$\begin{aligned}
\mathrm{E}(A_{k+1}) &= \mathrm{E}(\mathrm{E}(A_{k+1} \mid A_k)) \;=\; 2N\ \mathrm{E}((1-(1-1/N)^{A_k})) \\
\mathrm{var}(A_{k+1}) &= \mathrm{E}(\mathrm{var}(A_{k+1}|A_k)) \;+\; \mathrm{var}(\mathrm{E}(A_{k+1} \mid A_k)) \\
&= 2\ \mathrm{E}\left(N(1-1/N)^{A_k}(1-(1-1/N)^{A_k}) \;+\; N(N-1)\left((1-2/N)^{A_k} - (1-1/N)^{2A_k}\right)\right) \\
&\quad + 4N^2 \mathrm{var}(\mathrm{E}(1-1/N)^{A_k})
\end{aligned}$$

The overall cumulative means and variances are harder to obtain, since the distribution of $A_k$ is has no simple explicit form.

### B Extremes of survival and descent

This section continues the discussion of section 3.2.1 of survival and descent of a 30M genome in a diploid dioecious population of 100 males and 100 females at each generation. Here are presented results for the extreme individuals represented in Figures 12, 13 and 14.

Note the details of such extreme descent will be specific to the population pedigree, but each pedigree is likely to have a few individuals with extreme high or low descent patterns. For example, in the pedigree here, there is one diploid individual, $A$, who has 2 offspring, but then just 2 descendants for the subsequent 5 generations: he is the last founder to become ancestral to the entire population. The probability that an individual with 2 descendants eventually becomes ancestral to everyone is not surprising – the probability is about 95 %. However this particular trajectory of two descendants for 6 generations has probability only $1.8 \times 10^{-5}$ and the probability of existing in a population of 200 of only 0.004. On the other hand, the probability of a founder having descendants but no more than 2, in 6 consecutive generations (i.e. 1 or 2 each generation) is 0.0036, and the probability such an individual will occur in a population of 200 is just about 50%.

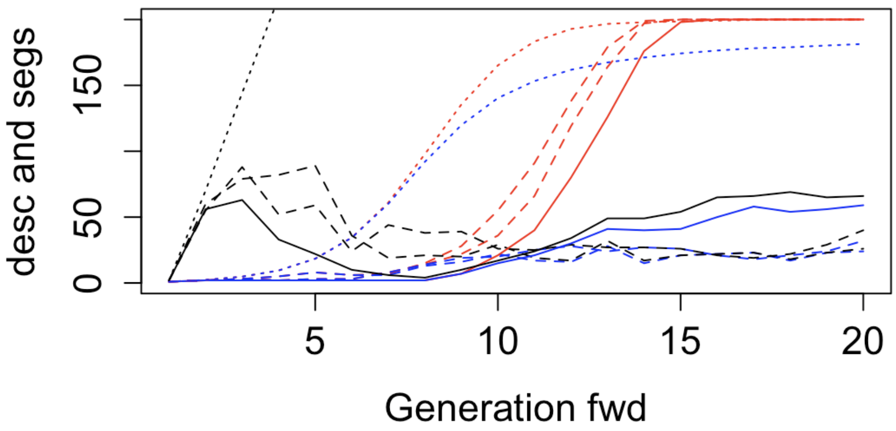


Figure B.1: Individuals with minimal descent in Figure 13 : for details see text

Figure B.1 shows more details of the individuals of Figure 13 with the lowest positive descent. Black lines show the number of descendant genome segments, red lines the number of diploid descendants, and blue lines the number of diploid genetic descendants. The dotted lines are the means over all individuals with positive descent, the dashed lines are the two individuals with the smallest positive number of genetic descendants at generation 40, and the solid lines are for the individual $A$ who is the last to become ancestral to all 200 individuals. Each blue line is bounded above by both black lines and red lines. The red lines are initially indistinguishable from the blue: for the first five generations all descendants/ancestors are typically genetic descendants/ancestors (Thompson, 2024).

Note that for all three extreme individuals by 20 generations the black line is not much greater than the blue: for these individuals most genetic descendants will carry only a single segment from the 30 Morgan genome. This is not in conflict with the results shown in Figure 15 of Section 3.2.2. While there is little surviving genome, and typically a single descendant segment in each genetic descendant, that same segment may be present in multiple descendants. By generation 5 these three individuals have descendant segments way below the mean, which already exceeds 200. One genome of $A$ is extinct and the other barely survives a "bottleneck" in descendants at generation 7 with a total of only 4 descendant segments of genome. However, $A$ thereafter becomes ancestral to everyone, and is a genetic ancestor of 25% of the population by generation 20. In contrast, the two individuals with the lowest final descent have more descendants initially, but lose significant lines of potential genetic descent at around 8 to 12 generations. As discussed in Thompson (2024), this is the period in which a large proportion of genetic ancestral lineages are lost. One genome of one of these two founders becomes extinct at generation 10. Both genomes of the other individual survive, but one has low segment count, including a 3-generation period from generation 27 to 29 in which there is only a single descendant segment from this genome.

At the other extreme are diploid individuals with maximal total descent at generation 40; there are three individuals with more than 250 Morgans of descent. They. have an average of 4.3 offspring, 10.7 grandchildren, and 22.7 great-grandchildren. This rapid initial expansion is reflected in the count of descendant segments. At generation 4 they have an average of 532.3 segments descending from their pair of 30M genomes, very close to twice the value of 216.80 for all founders with genome surviving at this generation. An alternative way to measure initial success is to consider the four founders who are first to become ancestors of the entire population. Four founders, including one of the previous three, achieve this at generation 9, after 8 generations of descent. (There are 25 others who achieve universal ancestry by the next generation.) These four founders have an average of 2.75 offspring, 9 grandchildren, and 21.5 great-grandchildren, showing a slightly smaller accumulating advantage over expectations: they are genetic ancestors of all these descendants. At generation 9 they are genetic ancestors of almost all, but not all, of their 200 descendants; 2, 8, 2 and 6 descendants are not genetic descendants. At generation 4, these four founders have an average of 480.5 descendant segments, smaller than for the previous 3 founders. However at generation 9 the 4 founders have an average of 1649 segments descending from their pair of 30M genomes. This is larger than the value of 1609 for the previous three founders, and may be compared with a mean of 582.6 for founders with genome surviving at this generation.

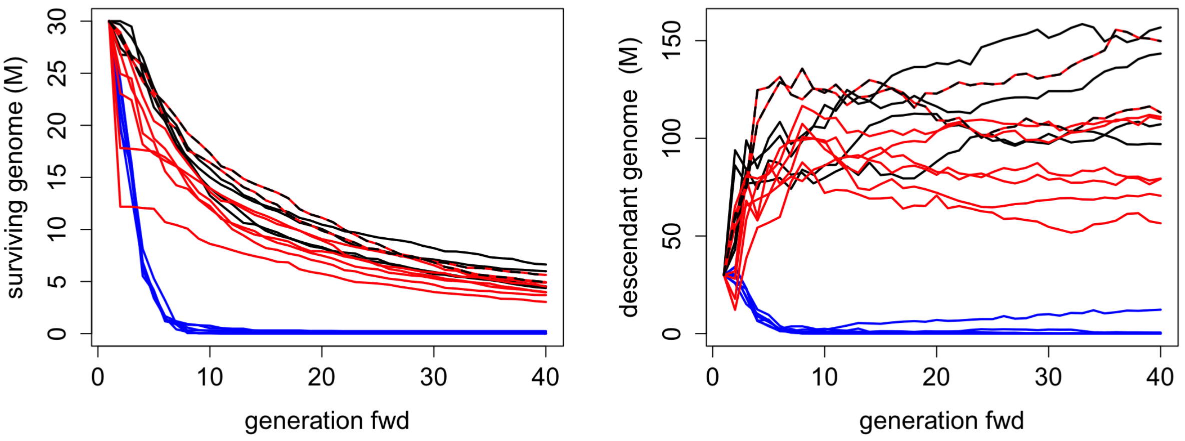


Figure B.2: Surviving and descendant lengths of genome for extreme individuals: for details see text

In Figure B.2 are shown the length of surviving genome, and of descendant genome (cf Figure 14)

for these three sets of individuals. In red are shown the value for the eight genomes of the four diploid individuals first to reach universal ancestry, and in black are the six genomes of the three diploid individuals who have over 250 Morgans descendant genome at generation 40. In blue are shown the values for the six genomes of the three diploid individuals with low descent: two of these six genomes in fact become rapidly extinct. The fates of founder genomes are largely determined by events that happen within the first five generations.

Six of the eight 'red' genomes have very similar trajectories of surviving genome; the other two correspond to an individual with only 1 offspring, but with 6 grandchildren. All these 8 genomes are present in 120 or more descendant genomes by generation 7. Thereafter the total frequency of the increasingly small descendant segments are determined by random genetic drift. Only one of the eight is among the maximally represented founder genomes at generation 40. The comparison of the 'black' and the 'red' genomes is of interest. The ultimate higher dominance of the 'black' genomes is due to their very rapid initial increase, which leads to not only higher ultimate descent, but also to a higher proportion of their genome surviving. In contrast the red genomes are particularly successful after generation 5, and thus spread more broadly in the population. Whereas the black genomes have substantially higher mean descent at generation 4, the red ones are higher at generation 9, the point at which they become ancestral to the entire population.

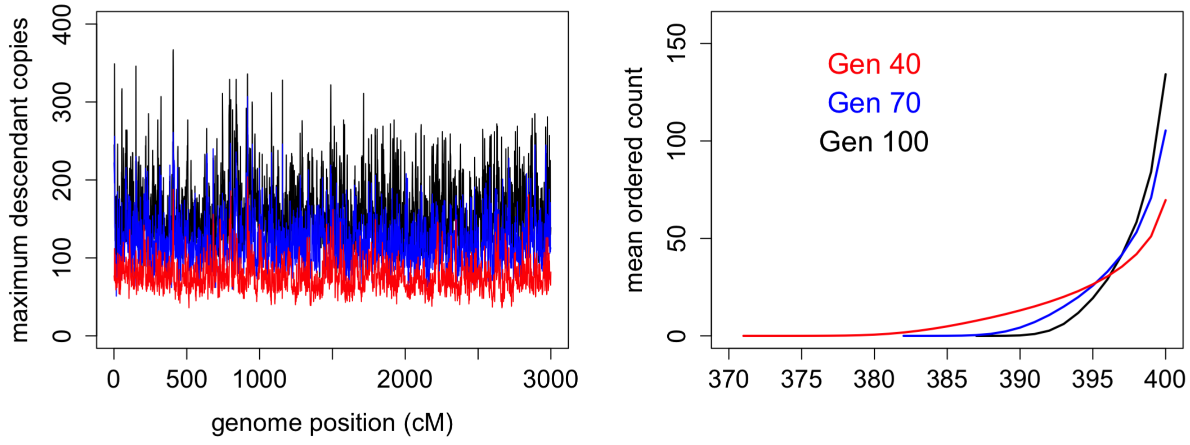


Figure B.3: FGL frequencies across the genome at 40, 70, and 100 generations. Left: The maximal frequency within each 1 cM of the 30M genome. Right; the mean across the genome of ordered FGL frequencies

In contrast the 3 individuals represented in blue have few descendants in the initial generations; surprisingly perhaps they all have two offspring, but only a mean of 3 great-grandchildren (generation 4). One genome of the individual $A$ who had only 2 descendants until generation 7 is extinct. The other recovers in terms of total descendant genome as the number of genetic descendants of the founder increases rapidly from 21 at generation 11 to 50 at generation 16. However these are replicates of the same small amount of surviving genomes: already by generation 7 only about 1% of this founder's genome survives, but is thereafter retained. In contrast the other two founders with low final descendant segments retain more surviving genome (about 2%) until generation 12, but by generation 40 all three have a very similar small amount of surviving genome.

Although there are individuals with a very large number of descendant segments with multiple segments present in all current individuals, interestingly there is no point of the genome at which any FGL is fixed. Figure B.3 shows on the left the maximum FGL frequency across the genome at 40, 70, and 100 generations, and on the right the average across the genome of the ordered FGL frequencies at these same time points. The maximum number of FGL surviving is 29, 18, and 13, respectively. Typically, by generation 100 there are about 7 or 8 FGL surviving. It is perhaps surprising that, in 30 Morgans and in a population of 400 haploids, no point of the genome coalescences to a single point by 100 generations (Kimura and Ohta, 1969; Greenbaum, 2015). It is a topic of future interest as to whether this is a chance outcome of this particular population pedigree, or whether it is perhaps a function of the diploid model which leads to very high correlation in survival and descent between the two FGL of each diploid founder (Section 3.4),

### C Survival and descent of 1-Morgan lengths of genomes

This section gives a number of Supplementary figures relating to Section 3.3, relating to the variation in descent and survival of a 1-Morgan length of genome..

#### C.1 Surviving genome within and among pedigrees

Figure C.1 shows for generations 5, 10, 15 and 20 the same results as presented for generation 10 in Figure 17. In Figure C.1 (Left) we see the declining amounts of surviving genome over the generations, and also the declining empirical standard deviations conditional on some genome surviving. On the right, we see the greatly decreased standard deviations, and increased count of diploid founders with extinct genomes, when instead of identifying founders we score the population distribution. See Section 3.3.1 for more details and discussion.

Likewise Figure C.2 shows for generations 5, 10, 15 and 20 the same results as presented for generation 10 in the corresponding Figure 18. We see the variation among population pedigrees (Right) barely exceeds the within-pedigree variation (Left). See Section 3.3.2 for more details and discussion.

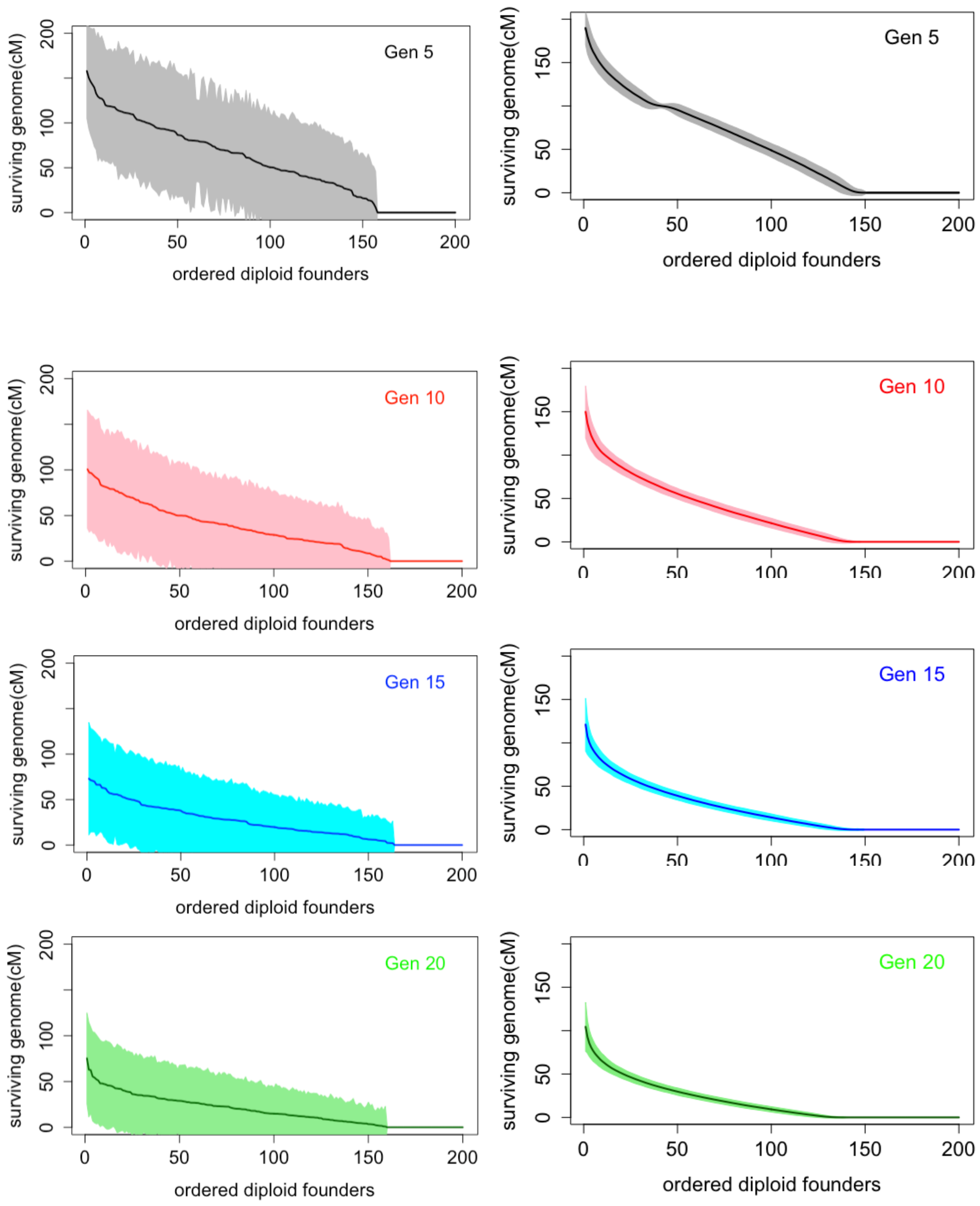


Figure C.1: Ordered survival amounts from 1-Morgan founder diploid genomes, with empirical standard deviations. Left: By founder individual. Right: By population distribution

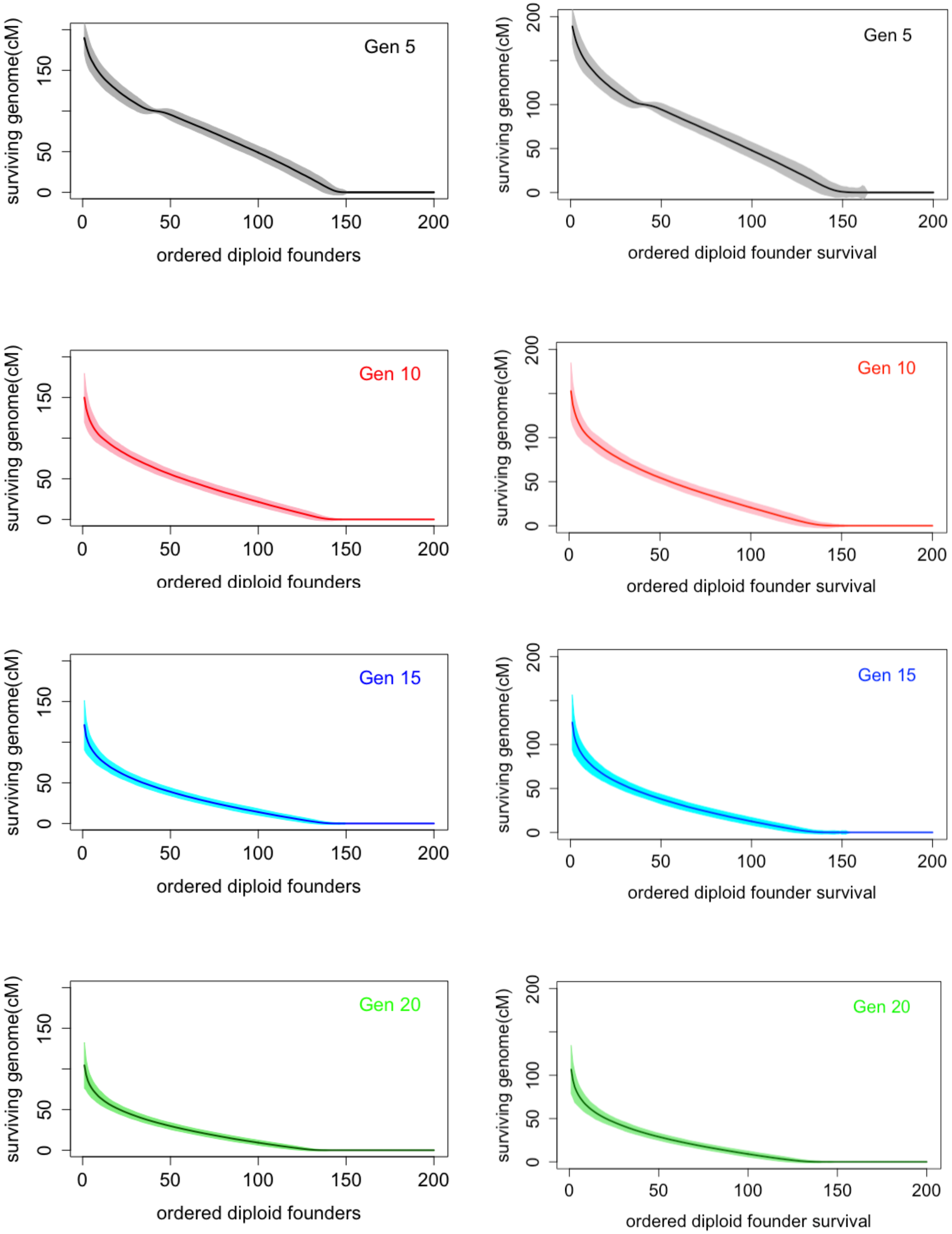


Figure C.2: Ordered survival amounts from 1-Morgan founder diploid genomes, with empirical standard deviations. Left: On a single population pedigree. Right: Over population pedigrees

#### C.2 The first few generations

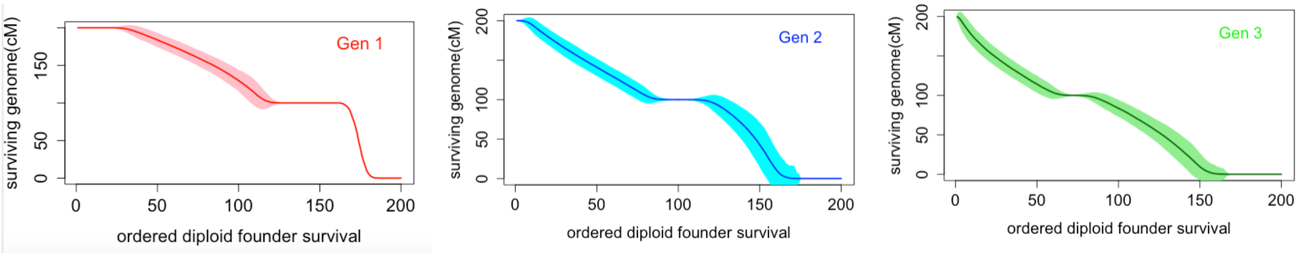


Figure C.3: Surviving genome in 500 random population pedigrees: the first few generations

In the population distributions of Figure C.2, at generation 5 there are always a few diploid founders who have exactly 100cM surviving. This is a relic of the very first meioses that disappears by generation 10. Figure C.3 shows the first three generations in detail, for the case of one realization on each of 500 population pedigrees. In the first generation, about 13.5% of diploids will have no offspring, and about 27% will have exactly one offspring. This offspring will receive exactly 100 cM from the founder parent, often but not always as a mosaic of the two founder genomes. There will be individuals who have one offspring in some realizations and 0 in others. The mean survival between 0 cM and 100 cM represents the proportion of realizations in which they have a single offspring, and conditional on having surviving genome there is no variation in the amount (100cM) of genome that survives. At the other end there are individuals with all their genome (200cM) surviving. The probability a 1-Morgan length is inherited intact is about 36%. There will be individuals with multiple offspring, who pass one haploid to one offspring, and the other to another; this is not the only way for the full 200 cM to survive in offspring, but it is the simplest and not rare. Individuals with more than one offspring, but passing a mosaic of their chromosomes, will have between 100cM and 200cM surviving, and only these will show variation in the surviving amount.

Generations 2 and 3 continue the pattern, and only-child Generation-1 individuals who pass an intact founder chromosome to Generation-2 will pass an exact 100cM of genome of some diploid founder parent. There will now be variation in survival amount less that 100 cM, more founder individuals will have no surviving genome, and even by generation 2 almost none have the full 200 cM surviving. Over generations 3 to 5 and beyond, as genome is lost the signature of an exact 100cM survival is also lost, but it is of interest that in a few percent of individuals it remains at 5 generations.

#### C.3 Maximal descendant counts over 1-Morgan lengths of genome

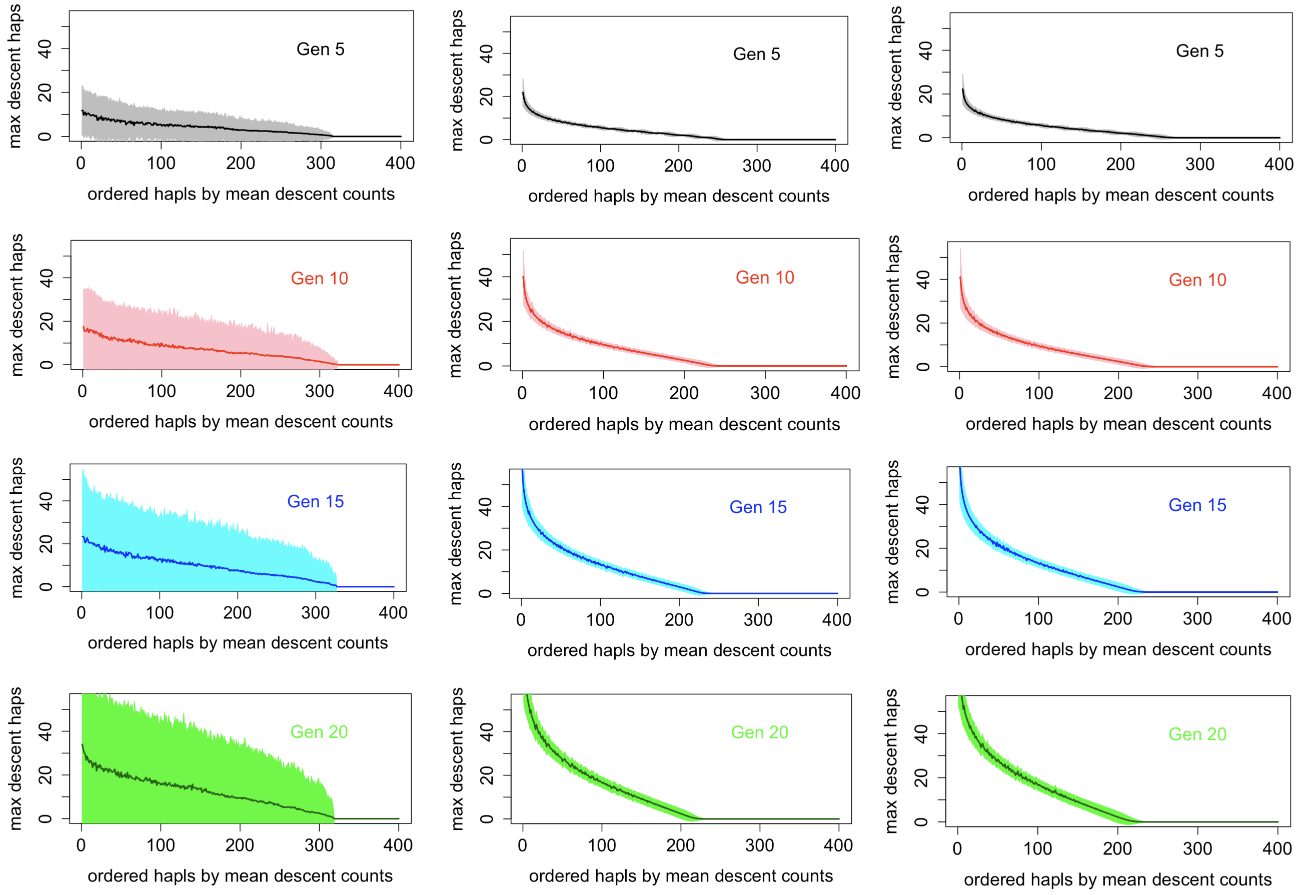


Figure C.4: Maximal descendant counts of founder haploids (FGL) across 1-Morgan. Left: Single pedigree by founder haploid. Center: Single pedigree by populations distribution. Right: Across pedigrees

Figure C.4 shows for generations 5, 10, 15 and 20 the same results as presented for generation 10 in the corresponding Figure 19 of Section 3.3.3. We see again the greatly reduced variance of the within-population distribution (Center) relative to individual variation (Left), both over realizations on the same population pedigree. We see also that variation over population pedigrees (Right) is barely greater that the within-population variance (Center). See Section 3.3.3 for more details and discussion.

### D Dependence in survival of founder genomes

The first two figures in this section relate to Section 3.4 and the study of dependence in descent in a population of 100 males and 100 females. The third figure related to the extension of this study to a larger population (Section 4.1.3).

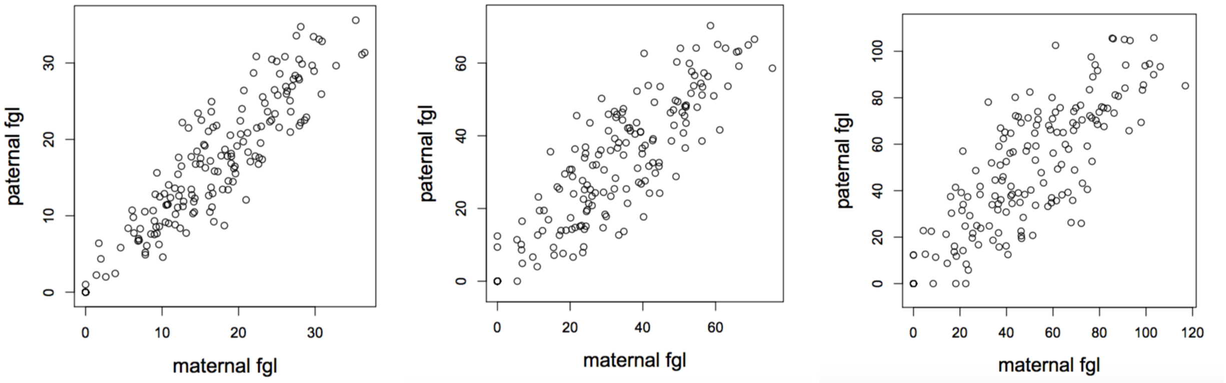


Figure D.1: Square root of number of descendant segments of a founder's maternal and paternal genomes in a random-mating population of 100 males and 100 females. Left: Generation 10. Center: Generation 40. Right: Generation 100.

Figure D.1 shows the corresponding scatterplots to Figure 20 showing dependence in descent between the two haploid genomes (FGL) of each founder. Here, instead of the counts of current genomes with genetic descent, the square root of the number of descendant genome segments is used at the measure for each FGL. We see that at the later generations this measure based on count of descendant segments gives less extreme marginal distributions, although the correlations (Table 1) are broadly similar.

Figure D.2 shows the same results for descent from either of the two FGL of founder mates. The upper row uses the count of current genomes containing any descent from an FGL (as Figure 20), while the lower row uses the square root of the number of descendant segments from each FGL (as Figure D.1 ). Positive correlations remain, but at a far lower level (see Table 1).

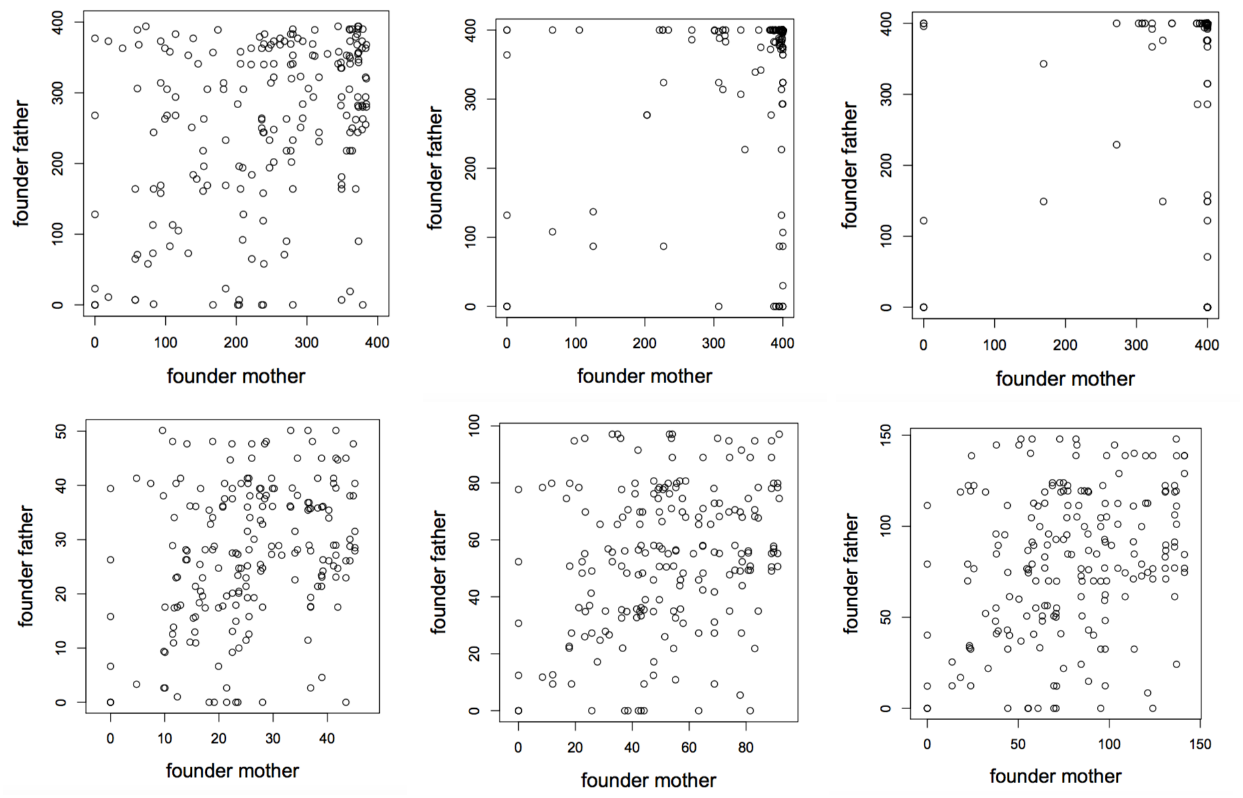


Figure D.2: Dependence in the descent of genomes of initial-generation mates in a random-mating population of 100 males and 100 females. Above: Number of current genomes having descent, Below: Square root of the number of descendant segments. Left: Generation 10. Center: Generation 40. Right: Generation 100.

Finally, we compare dependence in descent in a population of 100 males and 100 females (Section 3.4), with the same statistics in a larger populatiom of 1000 males and 1000 females (Section 4.1.3). Figure D.3 show the dependence in descent between the FGL of founders, and between FGL in founder mates. The correlations in these scatterplots are given in Table 2. For the smaller population the corresponding scatterplots are in Figures 20, D.1 and D.2 and the numerical correlations are in Table 1.

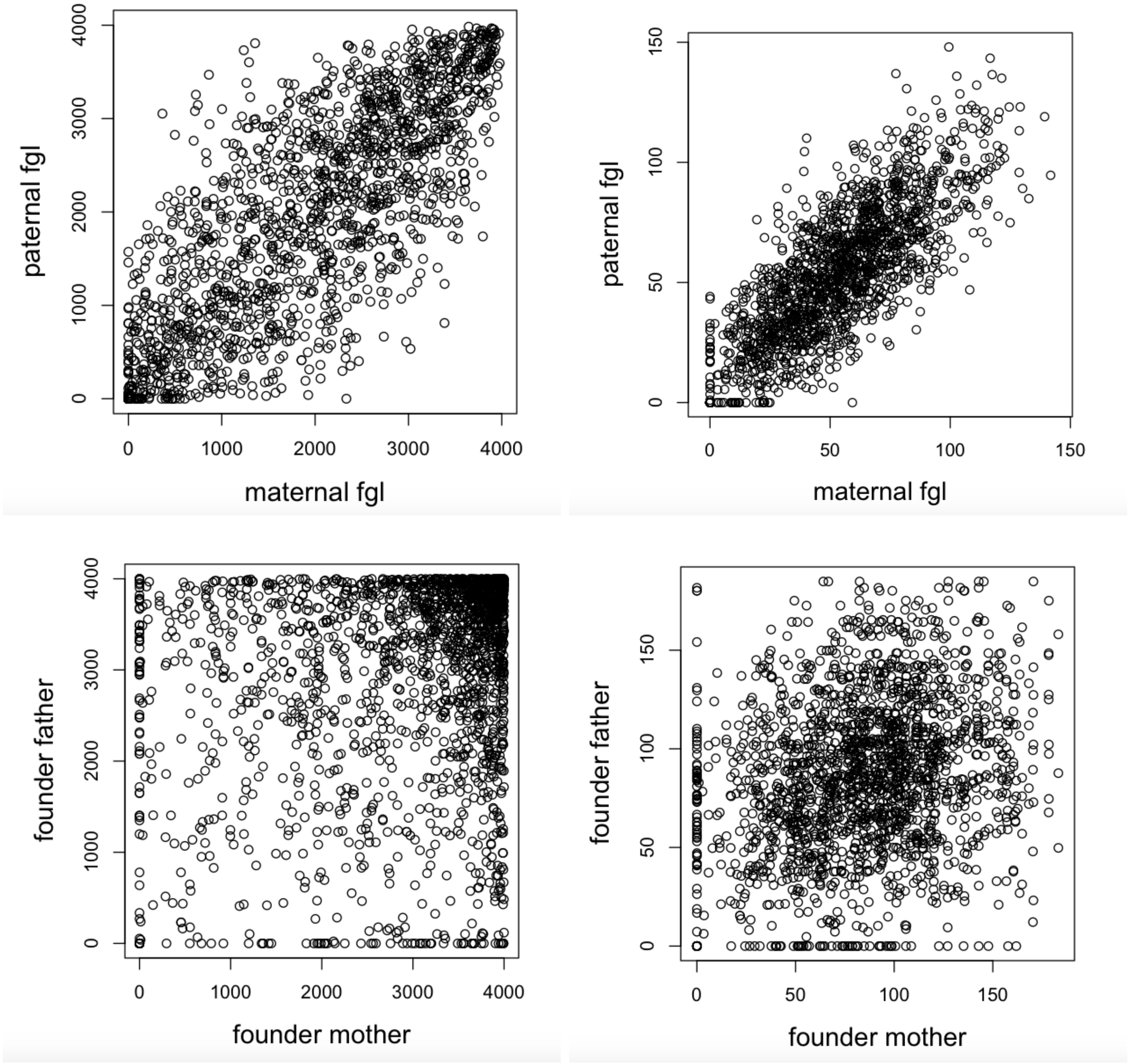


Figure D.3: Dependence in the descent of genomes in a population of 2000 individuals over 100 generations. Above: Dependence between the maternal and paternal FGL of each founder. Below: Dependence in descent from either FGL in pairs of individuals who are founder mates. Left: count of the final-generation descendant genomes carrying the specified FGL. Right: Square root of the total number of descendant segments from the specified FGL

### E When ancestors and descendants are equivalent

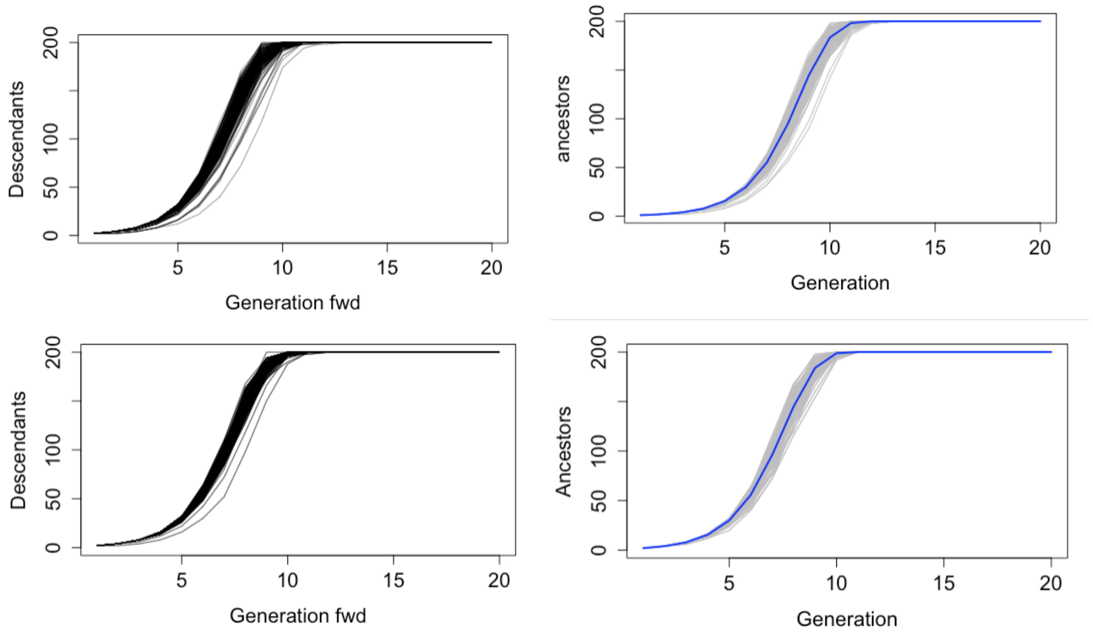


Figure E.1: Ancestors and descendants in a Zheng Cai population pedigree of 200 individuals per generation: Above: Over pedigrees. Below: Within pedigree. Left: Count of diploid descendants of each founder and Right: Count of diploid ancestors of each current diploid individual.

Zheng Cai, a visiting student (Pers comm, 2000), wished to simulate human population pedigrees, maintaining an equal number $N$ of males and females each generation. To achieve this she paired males and females together into $N$ couples, and gave each couple a son and a daughter. This is of interest here, as descendants and ancestors have the same count process. There is zero probability of extinction. Results are shown in Figure E.1. Ancestry of each current diploid converges quickly to all the diploid founders. Descendants of every diploid founder individual converges rapidly to the entire current population. There is minimal difference between population values (200 independent realizations), and pedigree values (the 200 individuals in a single population pedigree). As noted in the main text, convergence of ancestry/descent happens within 11 to 12 generations, a bit more quickly than the 16 to 17 generations for a random-mating population of this size.
